# Repetition-related gamma plasticity in macaque V1 and V2 is highly stimulus specific and robust to stimulus set size

**DOI:** 10.1101/2025.06.20.660699

**Authors:** Eleni Psarou, Mohsen Parto-Dezfouli, Iris Grothe, Alina Peter, Rasmus Roese, Pascal Fries

## Abstract

When a visual stimulus is repeated, the cortex has the opportunity to adjust its processing. Indeed, repeated stimuli induce reduced neuronal spike rates and increased neuronal gamma-band synchronization. Previous studies found the repetition-related gamma increase to occur both in human and non-human primates, for artificial and natural stimuli, to persist for minutes and to not transfer between strongly differing stimuli. Here, we further investigated the repetition-related effects using laminar recordings of multi-unit activity and local field potentials from awake macaque areas V1 and V2. We find that the effects on spike rate and gamma occur in all laminar compartments of V1 and V2. We quantify the degree of stimulus specificity with oriented gratings and find that the repetition-related gamma increase does not transfer to gratings differing by merely 10 °, the smallest difference tested. Furthermore, we find that the repetition-related effects are robust to stimulus set size, occurring both when one stimulus was repeated and when eighteen different interleaved stimuli were repeated. Finally, we show that alpha-beta activity increases and remains elevated when a stimulus is repeated, and decreases sharply when an unexpected stimulus is presented. These results suggest that repetition-related plasticity leads to changes in spike rates and rhythmic neuronal synchronization in different frequency bands that adjust the cortical processing of repeated stimuli.

## Introduction

Under natural conditions, we often stay in a given environment for some time. During such periods, surrounding stimuli typically do not change radically, and thus, the visual input we receive is constrained, i.e. not fully random, but repeating. Additionally, we often revisit with our eyes parts of the visual field that are of particular behavioral interest. For instance, while executing everyday tasks like preparing a cup of tea, we almost exclusively and repeatedly fixate task-relevant objects and parts of the visual field [1]. All this creates a considerable redundancy in our visual input, and the brain could build on this redundancy to optimize the processing of those repeated stimuli.

Previous studies have reported repetition-related plasticity in neuronal activity. Repeated stimuli have been shown to typically induce less spiking [2–5], and reduced hemodynamic responses [6, 7]. Interestingly, this repetition-related suppression is not linked to reduced behavioral performance. Instead, behavioral accuracy has been found to remain stable [8] or even improve with stimulus repetition, a phenomenon called ‘repetition priming’ [9, 10]. How could a significant decrease in spiking activity go hand in hand with sustained or even improved behavioral performance? In other words, what is the mechanism that allows the smaller number of spikes to maintain or even improve their downstream impact?

Increased synchronization at the neuronal network level has been proposed to allow fewer spikes to be transmitted in a more efficient way [11]. Indeed, several studies have linked stimulus repetition to increased synchronization [3, 4, 8, 12]. In particular, the repetition of visual stimuli has been linked to a profound increase in gamma-band activity within early and mid-level visual areas, as well as gamma-band synchronization between them, both in awake macaques [3, 4] and in human participants [8]. Between human visual areas, stimulus repetition predominantly increases bottom-up influences, consistent with increased or maintained information transfer. The abovementioned repetition-related changes have been shown to persist for several minutes [4, 8], and to be stimulus specific [4, 8], i.e. they do not transfer to other stimuli that have not been repeated.

So far, the stimulus specificity of the effects has been shown both for gratings [4, 8] and naturalistic stimuli [4]. These studies demonstrated that repetition-related plasticity did not transfer between gratings or natural stimuli that had been designed to differ strongly from each other. What happens when the repeated stimuli are more similar to each other? In this study, we systematically investigated the level and extent of stimulus specificity of the repetition-related plasticity. To do so, we used black-and-white gratings whose characteristics can be easily controlled and parameterized. By manipulating one dimension, the stimulus orientation, in fine steps, we were able to quantify the specificity of repetition-related plasticity. Importantly, all other stimulus characteristics remained unchanged. At the same time, this paradigm entailed blocks in which a larger number of different stimuli were repeated in an interleaved manner, whereas previous studies had interleaved maximally two to three stimuli [4]. This enabled us to test whether the repetition-related plasticity is also present under those conditions.

Furthermore, repetition is important in the context of theories on predictive coding. Those theories propose that feedforward circuits signal prediction errors, and feedback circuits signal predictions [13–15]. Feedforward signals emerge primarily from superficial layers of a given area and target granular layers of higher areas [16–18]; feedback signals arise (1) from deep layers and target deep and very superficial layers of lower areas, and (2) from superficial layers and target superficial layers of lower areas that are hierarchically close [19]. Thus, prediction/prediction-error related processes might show laminar specificity. Previous studies have shown that cortical rhythms also show a laminar pattern: alpha and beta is more prominent in deep layers [20, 21], while gamma is more prominent in superficial layers [20–23], though some studies report additional gamma peaks in Layer 4B and deep layers [22–25]. In agreement with this and the laminar origins of feedforward versus feedback projections, interareal directed influences in gamma are stronger in the feedforward than feedback direction, and influences in alpha/beta are stronger in the feedback than feedforward direction [26–30]. To investigate whether stimulus repetition leads to distinct changes between layers, we used the above-mentioned repetition paradigms while performing laminar recordings using linear probes in the monkey early visual cortex targeting areas V1 and V2.

## Materials and Methods

### Animals

Electrophysiological recordings were performed in one monkey. The rationale for using a single monkey is explained in more detail below, in the section on ‘sample size’. The monkey was implanted with a recording chamber over the occipital pole that allowed laminar access in areas V1 and V2. At the time of electrophysiological recordings, the monkey was 17 years old and weighed 15 kg. All procedures and housing conditions complied with the German and European law for the protection of animals (EU Directive 2010/63/EU for animal experiments). All experimental procedures were approved by the regional authority (Regierungspräsidium Darmstadt, Germany).

### Implants

The implant planning was based on pre-operative MRI (MAGNETOM Trio scanner, 3.0T; Siemens, Munich, Germany) and CT scans (Brilliance 6 scanner, Philips, Amsterdam, Netherlands; and ProMax 3D Mid scanner, Planmeca Oy, Helsinki, Finland) that allowed the extraction of the brain and skull models. First, the monkey was implanted with a cement-free headpost implant over the frontal brain areas [31]. A recording chamber was later implanted over the occipital pole. The location and angle of the recording chamber were planned according to the MRI scan. Special care was given so that the angle of the recording chamber allowed perpendicular access to the surface of V1 and thus, laminar cortical access. The bottom surface of the chamber was CNC-milled to follow the underlying skull geometry. A trepanation through the skull was performed 2.4 weeks after chamber implantation, which allowed access to brain areas V1 and V2. For the needs of another experiment (not reported here), a single virus injection was performed at one cortical location (at four different depths) within the trepanation. None of the recording sessions reported here overlapped with the injection site.

### Data collection Experimental setup

During the experiments, the monkey was head fixed and sat comfortably in a primate chair that was placed in a dimly lit Faraday cage that eliminated power-line noise and attenuated potential external sounds. All behavioral paradigms were designed and controlled by ARCADE, a stimulus presentation software written in Matlab (Dowdall, Schmiedt (32); https://github.com/esi-neuroscience/ARCADE). All stimuli were presented on an LG 32GK850G-B monitor (LG Electronics Inc., Seoul, South Korea.) with a resolution of 2560 x 1440 (16:9), a size of 697.34 x 392.256 mm and a refresh rate of 143 Hz. The following ‘default picture settings’ of the monitor were used: R/G/B: 50/50/50, Gamma Mode 2, Brightness 100, Contrast 70, Color Temperature Custom. The distance from the screen was 83.5 cm. Gamma correction was applied (gamma: 1.57). The exact timing of the stimuli was measured with a photodiode that was attached to one of the corners of the monitor (invisible to the monkey).

### Eye position monitoring

Eye signals of the left eye were recorded at 1 kHz using an Eyelink 1000 system (SR Research, Ottawa, Canada). The system was calibrated at the beginning of each recording session.

### Behavioral tasks

The stimuli used in the main experimental paradigm were circular patches of static black-and-white grating that always had the following features: diameter of 6.6 °, centered at 3 ° to the right of the vertical meridian and 3 ° below the horizontal meridian, spatial frequency of 2 cycles/°, contrast of 100%. Grating orientation varied from 0 ° to 170 ° in steps of 10 °. One orientation was presented per trial. For a given recording session, one orientation was randomly selected as the conditioned orientation and presented in a fixed-orientation block, in which this orientation was presented for 90 consecutive trials; this block is referred to as FIX. Before and after this conditioning, the orientation tuning of the recorded neuronal populations was assessed by presenting all employed orientations in a pseudo-randomly interleaved manner, with each orientation being presented 10 times; these trial blocks are referred to as variable-stimulus blocks, VAR1 and VAR2. This allowed us to compare the two orientation tunings, before and after conditioning, and quantify the specificity of conditioning-related changes.

In any given recording session, the monkey completed the following sequence of tasks: 1) Receptive field mapping, 2) Pre-conditioning orientation tuning (VAR1), 3) A block of approximately 10 minutes during which the monkey saw full-field flashes, referred to as “break” (intended for current-source density analysis, see below), and then could watch natural scenes or close his eyes, 4) Conditioning, i.e. repetition of the conditioned stimulus (FIX), 5) Post-conditioning orientation tuning (VAR2). In all tasks, the monkey was rewarded for maintaining central fixation throughout the trial without additional task requirements. For every correct trial, the monkey was rewarded with a drop of grape juice. While advancing the recording probe into the brain, the monkey occasionally performed additional blocks of full-field flashes.

### Receptive field mapping

The receptive fields (RFs) were measured using a black moving bar stimulus (bar width: 0.12 °, bar length: 15 °) that was presented on a grey background. In each RF-mapping trial, the bar was presented in one out of four orientations and was moved in one out of two possible directions, both orthogonal to the selected orientation. The bar moved at a speed of 8 °/s and covered the lower right visual quadrant, where the recorded RFs were expected, based on the known retinotopy of the recorded cortex. The monkey had to complete at least 80 correct trials, i.e. ten trials per condition.

### First and second variable-stimulus repetition block

The first and second variable-stimulus block were identical, and are referred to as VAR1 block and VAR2 block, or as VAR when referring to both. A grey background was presented throughout. Each trial started with the presentation of a white fixation point at the center of the screen. To perform a correct trial, the monkey had to turn his eyes towards the fixation point and maintain his gaze within a fixation window of 1-1.2 °radius around it throughout the trial.

Upon entering the fixation window, a baseline period of 0.8-0.9 s started. Then, an oriented grating was presented for 0.9 to 1 s. At the end of the stimulation period, the grating and the fixation point turned off, and a reward period of 0.8 s started during which the monkey received a drop of grape juice as reward. Then, an intertrial interval (ITI) of 0.5 s followed before the beginning of the next trial (Fig. 1A).

**Figure 1.**
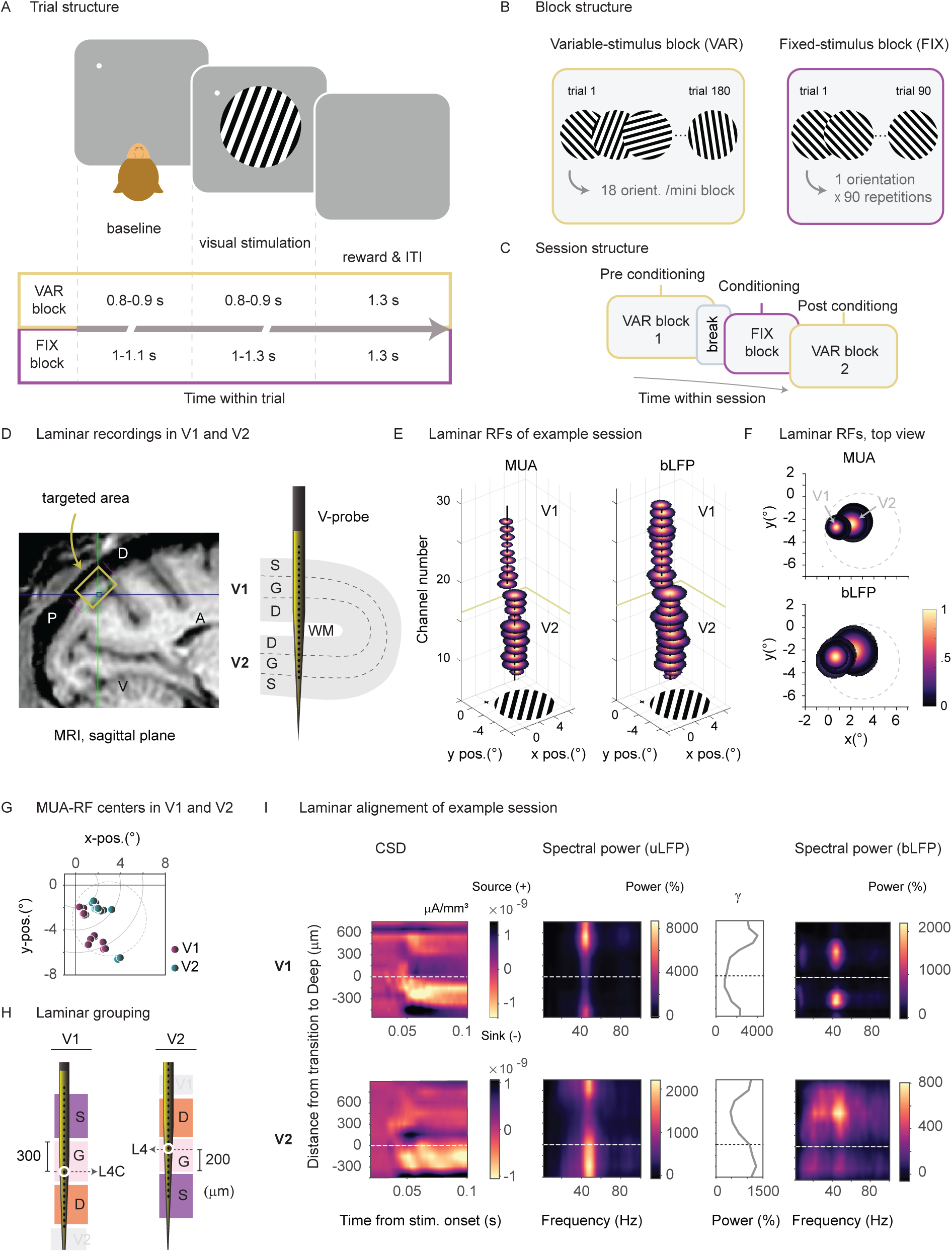
Behavioral task and electrophysiological recordings. (A) Schematic representation of the general trial structure and the specific timings for the VAR and the FIX blocks. (B) Structure of the variable-stimulus blocks, VAR1 and VAR2, and of the fixed-stimulus block, FIX. VAR blocks consisted of several mini-blocks. Within each mini-block, gratings of 18 different orientations were pseudo-randomly interleaved. In FIX, a stimulus with a fixed (for a given session), randomly selected orientation was presented across 90 consecutive trials. (C) Structure of the session, showing the sequence of blocks. Each session started with VAR1, which was followed by a short break and then by FIX. FIX was immediately followed by VAR2. (D) Left: Sagittal plane of an MRI scan showing the target area for electrophysiological recordings, marked by a yellow rectangle. The anterior (A) – posterior (P) and dorsal (D) – ventral (V) orientations are indicated. Right: Illustration of the targeted cortical area in V1 and V2 and their respective laminar compartments (S: superficial, G: granular, D: deep). Laminar recordings were performed with multi-contact linear arrays (V-probe). (E) MUA- and bLFP-Receptive Fields (RFs) in V1 and V2 of an example recording session as a function of channel number. The fixation spot (black cross) and the grating are shown on the bottom of the plots for comparison. (F) Top view of the RFs from (E). (G) MUA-RF centers per session, averaged over all contacts per recording session, separately for V1 (magenta, N=16 sessions) and V2 (dark green, N=16 sessions). The grey dotted line indicates the outline of the presented grating. (H) Illustration of the grouping of V1 and V2 recording sites into laminar compartments. (I) Neuronal responses to grating presentations in one example session each for V1 and V2. Shown from left to right: CSD, percent uLFP power change, percent uLFP-power change averaged over a gamma band frequency window (29-53 Hz), and percent bLFP power change relative to baseline. White dashed lines denote the transition from granular to deep layers (0 μm).

On any given trial, the orientation of the presented grating was randomly selected from a pool of 18 possible orientations (0 ° to 170 ° in steps of 10 °; Fig. 1B). When a given orientation was presented and the trial was successfully completed, that orientation was removed from the orientation pool. If the monkey broke fixation during a trial, the presented orientation remained in the pool. When all 18 orientations had been successfully presented, this set of trials constituted a completed mini-block. The pool was then refilled with all 18 orientations, and the procedure was repeated.

In VAR1, 10 mini-blocks were collected, so that 10 correct trials per orientation were acquired. In VAR2, at least 10 mini-blocks were collected, and then additional mini-blocks were collected until the monkey stopped working.

VAR2 followed immediately after FIX (Fig. 1C), i.e. there was only a regular ITI between the last trial of FIX and the first trial of VAR2.

### Fixed-stimulus repetition block

On each recording day, the monkey completed one fixed-stimulus block, referred to as FIX, during which a grating of a fixed orientation was repeated for 90 consecutive trials (Fig. 1B). At the beginning of this block, one orientation was randomly selected from the orientation pool (18 possible orientations; same as in VAR). We refer to this stimulus as the “conditioned” stimulus. The trial structure was identical to that of VAR, with the only differences that the baseline period was 1-1.1 s, and the stimulus period was 1-1.3 s. FIX ended when the monkey had successfully completed 90 correct trials. In case the monkey broke fixation during a trial, the trial was aborted and a period of 0.5 s was followed by the ITI (0.5 s). As mentioned above, the conditioning block was immediately followed by VAR2.

### Electrophysiological recordings

Laminar recordings were performed using linear multi-contact arrays. We used V-probes (Plexon Inc, Dallas, Texas, US) with 24 or 32 contacts and an inter-contact spacing of 100, 125, or 150 μm. All electrode arrays had a contact-site diameter of 15 μm and a tip-to-first contact distance of 300 μm. The diameter of the probes differed depending on their number of contacts. The 24-channel and 32-channel probes had a diameter of 210 and 260 μm, respectively. In our analysis, we pooled data from all recording probes. For more details see section ‘Laminar alignment of recording sessions’.

On any given recording day, a grid was added in the chamber and a laminar probe was advanced perpendicularly to the brain surface to target area V1. The probe was advanced through a guide tube using a precision hydraulic micromanipulator (MO-972A; Narishige, Tokyo, Japan). The guide tube featured a blunt tip that rested against the dura and provided extra stability to the recording probe. The probe was advanced with a speed of 10 μm/sec until the first few electrode contacts showed brain activity in the LFP signals. Then, the probe was advanced further at a lower speed (1 μm/sec) until neuronal activity was present on several recording contacts covering all V1 layers. A few of the most superficial contacts were intentionally kept outside of the brain to ensure coverage of the most superficial cortical layers.

When the target position of the probe was found, the probe was slightly retracted (for 500-1000 μm, with a speed of 1 μm/sec) in order to compensate for and help release potential tissue dimpling. This allowed us to approximately preserve the target recording position. A similar procedure has been described by Nandy, Nassi (33). Probe retraction was followed by a waiting period of approximately 30 minutes to allow the tissue to stabilize before recording. During this period, the monkey could watch videos of natural scenes or sleep.

### Electrophysiological signals

The recording ground was connected to the metal guide tube that protected the probe from bending and stayed outside of the brain with its flat tip resting firmly on the dura. The recording reference was connected to the shank of the linear probe.

Electrophysiological data were acquired with Tucker Davis Technologies systems (TDT, Alachua, Florida, United States). Raw data were recorded and digitized at 24414.0625 Hz using a TDT PZ2 preamplifier. Offline, the raw data were downsampled to 1000 Hz. For the LFP, this was achieved by first upsampling to 3,125,000 Hz, then downsampling to 25,000 Hz, applying an IIR filter with a stop-band at 500 Hz and downsampling to 1000 Hz. For the MUA, this was achieved by first upsampling to 3,125,000 Hz, band-pass filtering between 300 and 12000 Hz, rectification, IIR-filtering with a stop-band at 500 Hz and downsampling to 1000 Hz. This way of deriving MUA is similar to approaches used in several previous studies [4, 34, 35]. Note that there was no line-noise removal, because the recordings were done in a Faraday cage that completely avoided line noise.

### Data analysis

Data analysis was performed with MATLAB (Mathworks, Boston, USA) using the FieldTrip toolbox [36].

### Bipolar derivation

To reduce the influence of volume conduction on the LFP signals, local bipolar derivation was performed and used for all LFP analyses. The bipolar derivation was computed sample-by-sample as the value measured on a given electrode minus the value on the next-lower electrode, and we refer to a bipolar derivation as a (recording) site. We refer to the unipolar LFP as uLFP, and to the bipolar-derived LFP as bLFP.

### Receptive field analysis

RFs were estimated for each recording site, using similar approaches for MUA and bLFP signals. The MUA signal was low-pass filtered (Butterworth, order 2, backward and forward, low-cut at 100 Hz). For the bLFP signals, the time-resolved power at 40-100 Hz was estimated in time windows of 0.1 s length and in steps of 0.01 s. These neuronal signals were averaged over trials. Average MUA and bLFP responses were then baseline corrected and shifted in time according to the latency that gave the maximum response. We tested latencies between 0.04 and 0.07 s from stimulus onset, in steps of 0.005 s. The resulting responses were back-projected and plotted as activation maps [37].

### Estimation of receptive field centers

The centers of MUA and bLFP-RFs were estimated as follows. First, the RF maps were z-scored and then thresholded in order to eliminate noise. In particular, all RF-map elements below 5 SD were set to zero. The resulting RF maps were fitted with 2D-Gaussians (Fig. S1A). The coordinates of the Gaussian peak were then used to define the x and y coordinates of the RF center. Note the very good correspondence between MUA- and bLFP-RF centers (Fig. S1B). See Fig. 1G for an overview of all V1- and V2-RF centers of the recording sessions included in the analysis.

As shown in Fig. 1G, in two V2 sessions, the RF centers were located just outside of the grating stimulus. Note, however, that portions of these RFs still overlapped with the grating. Thus, these sessions were included in our analysis. Fig. S1C shows the MUA-RF centers of all sessions with simultaneous recordings in V1 and V2.

### Laminar alignment of recording sessions

On any given day, recording channels were first assigned to area V1, area V2, outside of the brain or within the white matter, based on their MUA- and bLFP-RF maps. The most superficial channels on the linear recording probes, which lacked clear RFs and showed less variability in the LFP signal, were considered to be located outside of the cortex. On some recording days, some of the deepest channels on the linear probe reached the underlying area V2. Channels were assigned to V2 when there was an abrupt shift in the eccentricity and an increase in the size of the RF compared to the RFs of the more superficial V1 channels (see example in Fig. 1E-F). Any deep channel (below V1 or between V1 and V2 channels) that lacked clear RFs was considered to be located in the white matter.

Then, V1 and V2 channels were further subdivided into three distinct laminar compartments: superficial, granular (input), or deep. The laminar assignment was informed by the CSD analysis, which allowed us to identify the input layer [38, 39].

Specifically, we estimated the CSD responses to the presentation of the gratings in the VAR and FIX blocks. We had originally planned to estimate the CSD based on the full-field flashes, but the CSDs based on the VAR or FIX blocks turned out to be more reliable, probably due to the larger number of trials. CSD analysis of LFP signals was performed using the MATLAB file exchange function ‘CSD’ [40]. The CSD of each channel was estimated per trial and then averaged across trials. For each electrode x, the CSD per trial was calculated using the following equation:

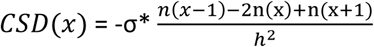

where, σ = 0.3 (S/m) is the conductivity of the extracellular medium, and h is the inter-contact spacing.

To ensure that there was no drift of the recording probe throughout the session, the CSD maps were calculated separately for VAR1, FIX and VAR2. Based on these separate CSD maps, we did not notice drifts of the earliest current sink, and therefore used all maps to inform our decision.

We first describe the procedure to assign electrodes to laminar compartments in V1. The electrode showing the earliest current sink was defined as the input layer 4C (L4C). On this electrode, the early current sink was consistently followed by a current source. The first or second electrode below the L4C electrode consistently showed a current sink that started after the L4C current sink [41, 42]. This electrode was defined as the uppermost electrode of the deep laminar compartment. The lowermost electrode of the deep laminar compartment was defined as the lowest electrode showing a clear bLFP-RF with V1 characteristics (see above). The granular compartment was defined to contain the L4C electrode and all electrodes up to 300 μm above L4C. Electrodes above this boundary were defined to be in the superficial laminar compartment (Fig. 1H), until the electrodes did not any more show bLFP-RFs.

The same procedure was applied to assign electrodes to layers in V2 while (1) taking into account that the laminar sequence is inverted (Fig. 1H), (2) defining the granular compartment to be only 200 μm thick (instead of 300 μm in V1, Ziemba, Perez (43), Kelly and Hawken (44)). In three recording sessions, we obtained only a small coverage of area V2. In these cases, channels covering the first 300 µm were considered part of the deep layers of V2.

### Data inclusion

Our analysis included only behaviorally correct trials. Recording sessions that lacked a clear early current sink were excluded from our laminar analysis. In the case of area V1, sessions that yielded few MUA RFs, such that the MUA RFs spanned < 800 μm of cortical tissue, were excluded from the analysis.

After application of these selection criteria, 16 V1 sessions remained from a total of 22 sessions that targeted V1, and 16 V2 sessions remained from a total of 16 sessions that targeted V2. Seven channels from 3 sessions were excluded due to the presence of strong artefacts in multiple trials.

The quality of spiking activity showed a large variation between sessions and channels. To ensure that only channels with clear visually induced spiking responses were included in our analysis, we performed the following procedure: For each session and channel, a sign test was performed on the data from the first variable-stimulus block to assess if there is a significant increase in the spiking activity around the onset of the visual stimulus. Only channels that showed a significant increase in their spiking activity between the baseline (-0.6 to 0 s) and the stimulus presentation period (0 to 0.6 s) were included in further analysis of spiking activity. The number of channels included in the analysis is summarized in Table 1.

**Table 1.**
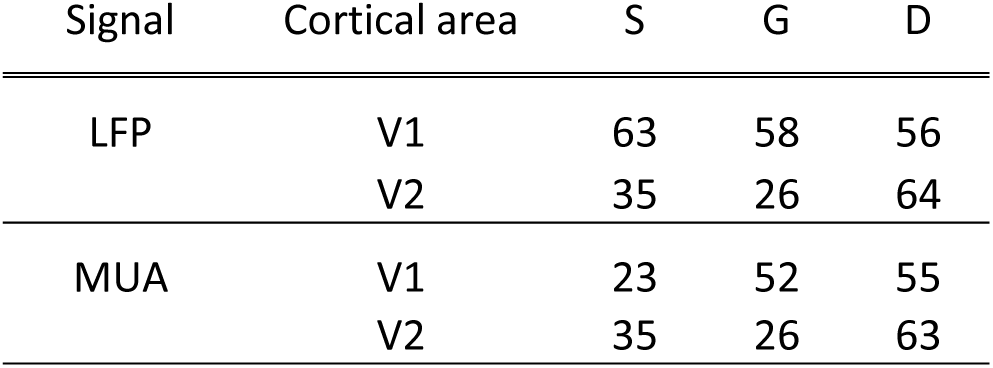
Number of included channels in LFP and MUA analyses per cortical area and laminar compartment (S: Superficial, G: Granular, D: Deep).

| Signal | Cortical area | S | G | D |
| --- | --- | --- | --- | --- |
| LFP | V1 | 63 | 58 | 56 |
|  | V2 | 35 | 26 | 64 |
| MUA | V1 | 23 | 52 | 55 |
|  | V2 | 35 | 26 | 63 |

In the analysis of intra-areal GC, we noticed that one session produced outlier results for V2. Specifically, when we calculated the mean and SD across the GC values from all V2 sessions, this session exceeded the mean plus 4 SD, and we therefore excluded it from the GC analysis.

### Spectral analysis of sustained period

We performed spectral analyses of bLFP power, and bLFP-bLFP GC. For all those spectral analyses, we used a 500 ms stimulus period from 270 to 770 ms post stimulus onset. This stimulus period was chosen to focus the spectral analysis on the sustained period and avoid the early post-stimulus onset transients.

All spectral analyses used the Fourier transforms of the bLFP signals in the described periods. Each 500 ms period was cut into five overlapping 300 ms epochs (Welch’s method), the epochs were linearly detrended, Hann tapered, zero-padded to 1 s, and Fourier transformed.

Power spectra were derived by squaring Fourier spectra. The power spectra of the visual stimulation period were baseline corrected. An average baseline power spectrum was estimated by combining all correct trials from all three repetition blocks over a pre-stimulus baseline window of 500 ms (-530 to - 30 ms relative to stimulus onset). The stimulus-period power spectra were then expressed as percent change relative to the baseline power spectra.

To define frequency windows of interest for both V1 and V2, we first averaged the power spectra over all V1 and V2 sites and all correct trials of FIX. The average spectrum showed clear spectral peaks at 9, 20, 41 and 80 Hz (Fig. S2). Correspondingly, we defined an alpha rhythm peaking at 9 Hz, a beta rhythm peaking at 20 Hz and a gamma rhythm peaking at 41 Hz. We did not consider a second gamma rhythm, because it was at a harmonic frequency and much weaker than the main gamma peak. The range of each frequency band was defined as +/- 30% around these peak frequencies, resulting in an alpha band of 6-12 Hz, a beta band of 14-26 Hz, and a gamma band of 29 to 53 Hz as indicated in Fig. S2.

GC spectra were derived through non-parametric spectral matrix factorization as described in [45] and implemented in FieldTrip [36].

### Single-trial estimates

To obtain single-trial estimates of MUA responses (stMUA), we defined a stimulus period (0.05 to 0.1 s relative to stimulus onset). Note that this stimulus period was meant to capture the strong transient MUA response and was different from the stimulus period for spectral analyses. The stMUA estimates were baseline normalized and expressed as percent change relative to baseline.

Single-trial estimates of power changes relative to baseline were obtained by averaging the power changes over the frequency windows of interest described above. Single-trial estimates for the gamma rhythm are referred to as stGamma. In some of the analyses, alpha and beta rhythms showed qualitatively the same effects (see Results) and therefore were combined into a common alpha-beta rhythm with a frequency band of 6 to 25 Hz (stAlpha-Beta). The single-trial power changes were baseline normalized as described in section ‘Spectral analysis of sustained period’.

### Assessing the repetition effect as a function of orientation difference from the conditioned stimulus

For a given recording session, the conditioned orientation was randomly selected from a pool of 18 orientations. We aimed at investigating the effect of repeating this orientation 90 times during the FIX block on the response to all 18 orientations. Therefore, we compared the responses during VAR2, obtained after the conditioning, to the responses during VAR1, obtained before conditioning. As VAR1 and VAR2 contained all 18 orientations, we could express the orientation presented in each trial as the orientation difference from that session’s conditioned orientation (i.e. the orientation presented in FIX). Positive and negative orientation differences of the same magnitude were combined (because there was no reason to assume that they were different), leading to orientation differences from 0 to 90 °. Note that in the VAR1 block, the monkey completed 10 mini-blocks, whereas in the VAR2 block, the monkey typically completed more mini-blocks. For this analysis, only the first ten mini-blocks of VAR2 were included to avoid potential biases due to different amounts of data.

We intended to estimate the neuronal responses just before and just after the FIX block, separately for each orientation and thereby for each orientation difference relative to the conditioned orientation (see Fig. 6A for an illustration). Each orientation (difference) was presented once per mini-block, and thereby 10 times during VAR1 and 10 times during VAR2. These repetitions inside the VAR blocks themselves were expected to incur repetition-related changes in neuronal responses, which we aimed at eliminating. Therefore, we performed regression analyses of the neuronal responses as a function of these 10 repetitions, separately per orientation difference, and separately for the VAR1 block and the VAR2 block. From the VAR1 regression, we obtained the estimate of the neuronal response at the end of the VAR1 block, just before conditioning and refer to it as the pre-conditioning value; From the VAR2 regression, we obtained the estimate of the neuronal response at the beginning of the VAR2 block, just after conditioning and refer to it as the post-conditioning value. The difference between the post- and the pre-conditioning value isolates the effect of the conditioning during the FIX block, while eliminating or minimizing effects of stimulus repetitions inside the VAR blocks (which were required to obtain sufficient data for robust estimation and statistics).

### Assessing the repetition effect as a function of the orientation difference from the preferred orientation of the recording site

We also analyzed the repetition effect as a function of the difference between the stimulus orientation and the preferred orientation of the recording site. The approach for this analysis was identical to the approach described in the last paragraph, except that the orientation difference was not relative to the conditioned orientation but relative to the preferred orientation of the recording site. The preferred orientation was determined as described in Womelsdorf, Lima (46): Let r_m_ (m = 1; 2; . . . ; 18) be the empirically observed neuronal response when the m^th^ stimulus orientation was presented, with the stimulus orientation denoted as θ_m_ = (0, 10, 20, 30,…, 180 °). We intended to express neuronal responses as complex vectors and perform a complex vector average, to use the resultant vector orientation as the estimate of the preferred orientation. However, stimulus orientation is a circular variable in the interval [0, 180]. Therefore, to use this approach, we scaled the orientation variable to a circular variable in the interval [0, 360] and transformed it to radians. For every m, we defined

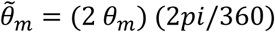

The vector sum was then defined as

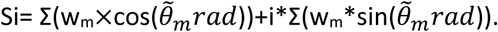

where w_m_ is the mean LFP response value of the m^th^ stimulus orientation. Subsequently, the orientation of the vector sum Si was extracted based on the inverse tangent, and it was re-scaled from the [–π, +π] interval to the [0, +π] interval, and re-transformed from radiant to degree. Finally, the nearest stimulus orientation to the orientation of the vector sum was considered as the preferred orientation. This procedure was applied based on the responses measured during VAR1, separately per recording site.

### Statistical analysis

All statistical analyses were based on non-parametric randomization tests.

We first describe the analyses for Figs. 2-5, which are based on two types of tests: (1) We tested for significant differences between repetition groups 1 and 3, as defined in the results and Fig. 2A; (2) We tested for significant slopes as a function of repetition number. Approach (2) takes all repetitions (except the early repetitions of FIX) into account, yet requires an estimation of the dependent variable for each single trial, which is difficult for GC. Therefore, we complemented it with approach (1), which defined trial groups that allow the straightforward estimation of GC; as this requires relatively many trials, we performed it only for the fixed-stimulus repetition block, containing 90 repetitions of the same stimulus.

**Figure 2.**
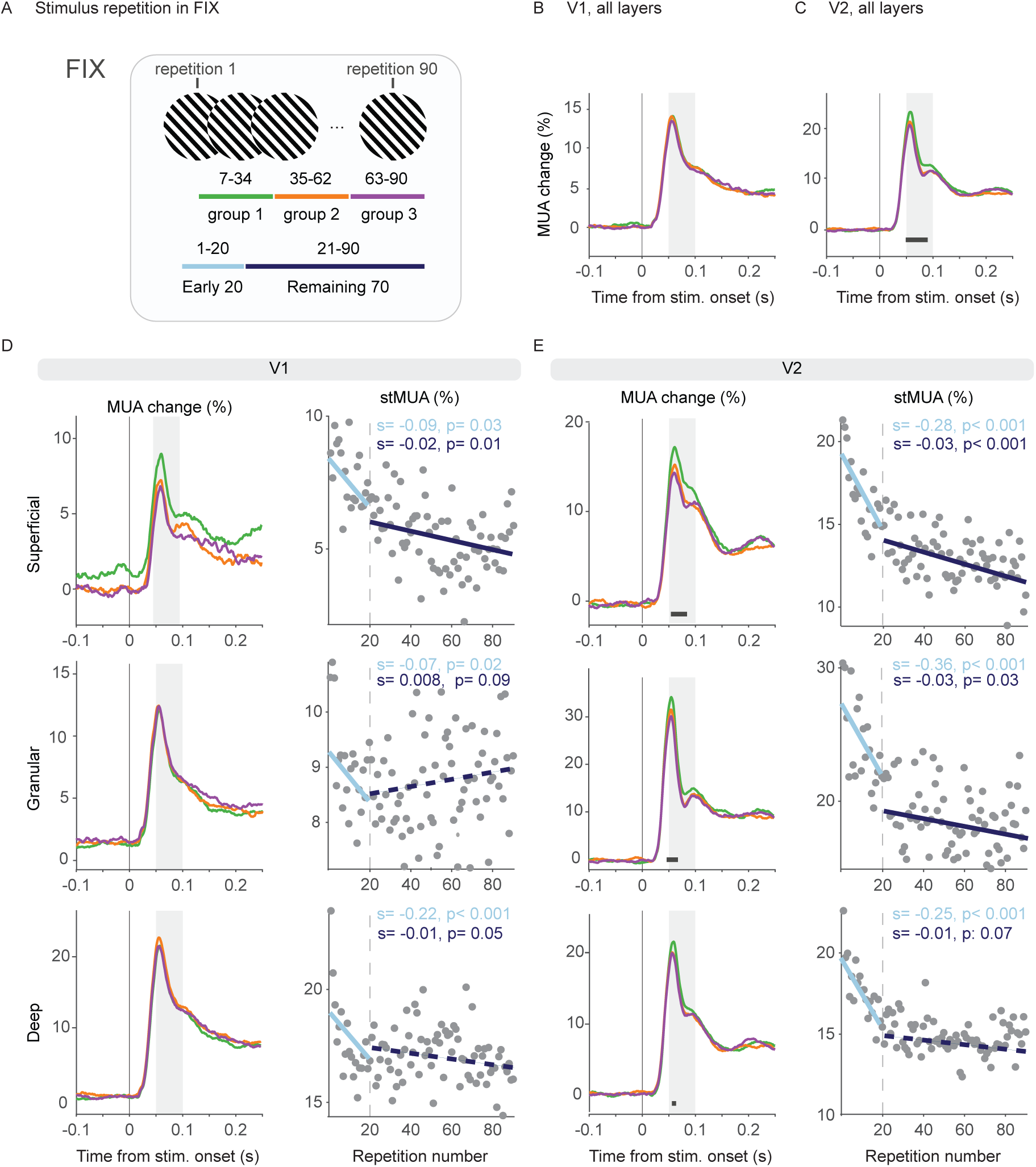
Repetition-related changes in MUA during FIX. (A) Schematic representation of the stimulus repetitions in FIX and their grouping into repetition groups 1, 2 and 3. Each group consisted of 28 repetitions. For the analysis of stMUA in D and E, we also defined an Early-20 and a Remaining-70 trials group. (B-C) MUA response from baseline averaged over all sites in V1 (B) or V2 (C), per repetition group (color-coded as in A). (D) Left: MUA response from baseline per repetition group averaged per laminar compartment (from top to bottom: superficial, granular, and deep). Grey rectangles indicate the time period that was used to estimate the stMUA responses. Horizontal black bars indicate periods of significant difference between groups 1 and 3. Right: stMUA as a function of repetition number within FIX. The dashed grey vertical lines indicate the 20^th^ repetition. Two separate regression fits are plotted: one for the Early-20 repetition group and one for the Remaining-70 repetition group. Regressions with (non-)significant slopes are plotted as (dashed) solid lines. The respective slopes (s) and p-values (p) are reported on top of each panel (see Methods for details). E) Same as in D but for V2. Note that y-axes were scaled per plot to optimally illustrate the effects.

Approach (1) compared spectra or time courses and therefore involved correction for multiple comparisons across frequencies or times, respectively. Per session and electrode or site (pair), we calculated differences between repetition groups 1 and 3 and subsequently averaged those differences over electrode or site (pairs) and sessions. We then performed 1000 randomizations. In each randomization, per session and electrode or site (pair), we randomly exchanged trials between groups 1 and 3 and re-calculated the difference. We then averaged over electrodes, sites or site pairs. We then retained the minimal difference and placed it into the distribution of minimal randomization differences, and we retained the maximal difference and placed it into the distribution of maximal randomization differences. The 2.5^th^ percentile of the minimal randomization distribution and the 97.5^th^ percentile of the maximal randomization distribution were used as thresholds for statistical significance, corresponding to a two-sided false-positive rate of 5%, corrected for multiple comparisons [47].

This approach allowed us to test for effects over time in the MUA, and over frequency in the case of power and GC spectra. In the case of GC spectra, we complemented our analysis with an additional statistical assessment (see motivation in Results) that looked for significant changes in three specific frequency bands: alpha (6-12 Hz), beta (14-26 Hz) and gamma (29-53 Hz) (see section ‘spectral analysis of sustained period’ above for the definition of these frequency bands). This approach compared the GC for three frequency bands and two directions, and therefore involved correction for six multiple comparisons. Per session and site pair, we calculated differences in GC between repetition groups 1 and 3 and subsequently averaged those differences over the frequencies within each frequency bands, separately per band, and subsequently over site pairs and sessions. This delivered one observed difference per frequency band and GC direction. We then performed 1000 randomizations. In each randomization, per session and site pair, we randomly exchanged trials between groups 1 and 3, re-calculated the difference and the averaging as for the observed differences, and kept the maximal and minimal differences separately per frequency band and GC direction. This resulted in one maximal randomization and one minimal randomization distribution per frequency band and per GC direction. Note that this approach was motivated by the large differences in GC values and GC differences between frequency bands, as discussed in the Results section. From each randomization distribution, we used the 2.5/6 percentile and the 100-(2.5/6) percentile as thresholds for statistical significance of the corresponding observed difference. Thus, the correction for multiple comparisons across the six combinations of frequency bands and GC direction used the Bonferroni method. This constitutes a non-parametric significance test with a two-sided false-positive rate of 5%, corrected for multiple comparisons across frequency bands and GC directions.

Approach (2) tested the slopes for single-trial values of MUA in a specific time window or spectral power in specific bands and therefore did not involve correction for multiple comparisons across frequencies or times. First, the data were averaged over sites, and subsequently a linear regression was performed to obtain the slopes for the observed data. For the VAR blocks, this gave separate slopes per session and per stimulus orientation, which were then averaged; for the fixed-stimulus block, this gave separate slopes per session, which were then averaged. Then, we performed 1000 randomizations. In each randomization, the repetition order was randomized, and subsequently, the same analyses were performed as for the observed data. The resulting slopes were placed into the randomization distribution. The 2.5^th^ and 97.5^th^ percentiles of this distribution were then used as thresholds for statistical significance with a two-sided false-positive rate of 5%. Note that some slope analyses were performed for specific repetition ranges, e.g. for the first 20 trials, which is then specified in the results and figure legends.

For Fig. 6, we used the difference between the above described post-conditioning value and the pre-conditioning value. Note that the randomization between VAR1 and VAR2 was performed without keeping the order within the respective blocks. This automatically eliminated the repetition effects expected to occur within the VAR blocks. Thereby, we could use, for the randomized data, the same test statistic as for the observed data, namely the difference between the neuronal response at the beginning of VAR2 minus the neuronal response at the end of VAR1.

For Fig. 7, we tested for significant differences in the mean neuronal responses between the three blocks (VAR1, FIX and VAR2). We compared the results of the regression analyses of their single-trial estimates in the following way. First, the mean (over sites and sessions) values of the fitted regression lines were computed for each block individually and subtracted from each other in order to obtain the observed mean difference between blocks. Then, we performed 1000 randomizations by shuffling single-trial estimates between blocks and repeating the analysis. This resulted in a distribution of mean differences, and its 2.5^th^ and 97.5% percentiles were used to as thresholds for statistical significance, as described above.

### Sample size

This study involved one male macaque monkey (*Macaca mulatta*). The use of a single subject allows an inference on that sample, but not on the population. If a second macaque had been added, any useful inference would still have pertained to that sample of two macaques, but not to the population of macaques, as we have shown previously [48]. As the inference remains qualitatively the same, we opted for a single macaque to reduce the number of animals used in research, according to the 3R principles [49]. This approach has recently been questioned [50], and we responded to this, as described in Psarou, Katsanevaki (51).

## Results

Laminar recordings of multi-unit activity (MUA) and local field potentials (LFPs) were obtained from primary visual cortex, V1, and in a subset of sessions from secondary visual cortex, V2, in one awake macaque monkey (Fig. 1D). The monkey performed a fixation task, during which large grating stimuli were presented on a screen; in each trial, one grating orientation was shown (Fig. 1A). In each session, the monkey completed the following trial blocks (Fig. 1B-C): (1) A first block of stimulus repetitions, where the grating orientation could vary from trial to trial, referred to as the “VAR1” block; (2) A short break; (3) A block of stimulus repetitions, where the grating orientation was fixed, referred to as “FIX” block; (4) A second block of stimulus repetitions with variable grating orientations, referred to as the “VAR2” block. This design allowed us to investigate neuronal responses for all stimulus orientations during the VAR blocks. Thereby, we could investigate the stimulus selectivity of the neuronal response changes between VAR2 and VAR1, which were induced by the repetition of the conditioned stimulus during FIX. Stimuli were stationary square-wave gratings of 18 different, evenly spaced orientations (see Methods for details). VAR1 contained 10 mini-blocks, with each mini-block containing one correctly performed trial for each of the 18 different orientations. VAR2 had the same structure as VAR1, and contained at least 10 mini-blocks, but then continued for as long as the monkey kept performing the task. FIX contained 90 correctly performed trials for one of the orientations, which was randomly selected from the 18 orientations for a given recording session; this orientation will be referred to as the “conditioned” orientation.

As shown in Figure. 1D-E, the recording chamber allowed perpendicular access in V1 and the underlying V2. Using linear multi-contact probes, we were able to simultaneously record across depths in both areas, yet not always across all depths in both areas simultaneously. Figure. 1E shows the receptive fields (RFs) of an example recording session. Within each area, the RF centers were aligned across depths, confirming that the recordings were mainly confined within the same cortical column. An abrupt shift of the RF centers, accompanied by an increase of the RF size indicated the transition from area V1 to V2 (Fig. 1E-1F; see also Fig. S1D). Figure 1G and Figure S1B provide a summary of the MUA- and bLFP-RF centers of the recorded sessions. The laminar recordings enabled the investigation of the previously described repetition-related changes as a function of cortical layers.

The VAR1 block allowed to investigate whether the repetition-related changes occur during the interleaved presentation of 18, i.e. many, different orientations, with a given orientation repeating only across sub-blocks, i.e. typically spaced by several (on average 17) intervening stimuli. This block also provided orientation tuning curves. The FIX block induced repetition-related plasticity for the conditioned orientation. The VAR2 block provided orientation tuning curves after the repetition of the conditioned orientation in the FIX block. The comparison of VAR2 with VAR1 allowed us to investigate whether changes induced by the repetition of the conditioned orientation during FIX affect neuronal responses induced by other orientations, and therefore, test how orientation-specific these changes are. Note that the conditioned orientation in the FIX block was randomly selected on each recording day. This dissociated the conditioned orientation from the preferred orientation of the recorded neuronal populations.

We set out to investigate stimulus-repetition effects as a function of cortical depth. To this end, we assigned for each session the recording electrodes to three laminar compartments: superficial, granular and, deep. We first computed the current source density (CSD) of the visually evoked LFP (Fig. 1I). Based on the CSD maps, we identified the input layers of areas V1 and V2. This allowed us to estimate the distance of each recording site from the input layer. This relative distance was then used to assign each recording site to one of the three laminar compartments (see Methods for a detailed description). Figure 1I shows the CSD and spectral power as a function of cortical depth of example V1 and V2 sessions.

The presentation of the grating induced a clear MUA response in all laminar compartments of both areas (Fig. 2D-E). The visual stimulation also induced a clear narrow-band gamma peak during the period of sustained stimulation (0.27-0.77 ms relative to stimulus onset) in all compartments of V1 and V2 (Fig. 3D-E). Additional spectral peaks were evident in both areas. In all V1 layers, there was a clear alpha peak centered around 8-10 Hz. An additional beta peak was present at 18 Hz in superficial and deep layers. In V2, there was a large peak in the beta range (20-23 Hz) and a smaller alpha peak (9 Hz) that was mostly evident in superficial and deep layers.

**Figure 3.**
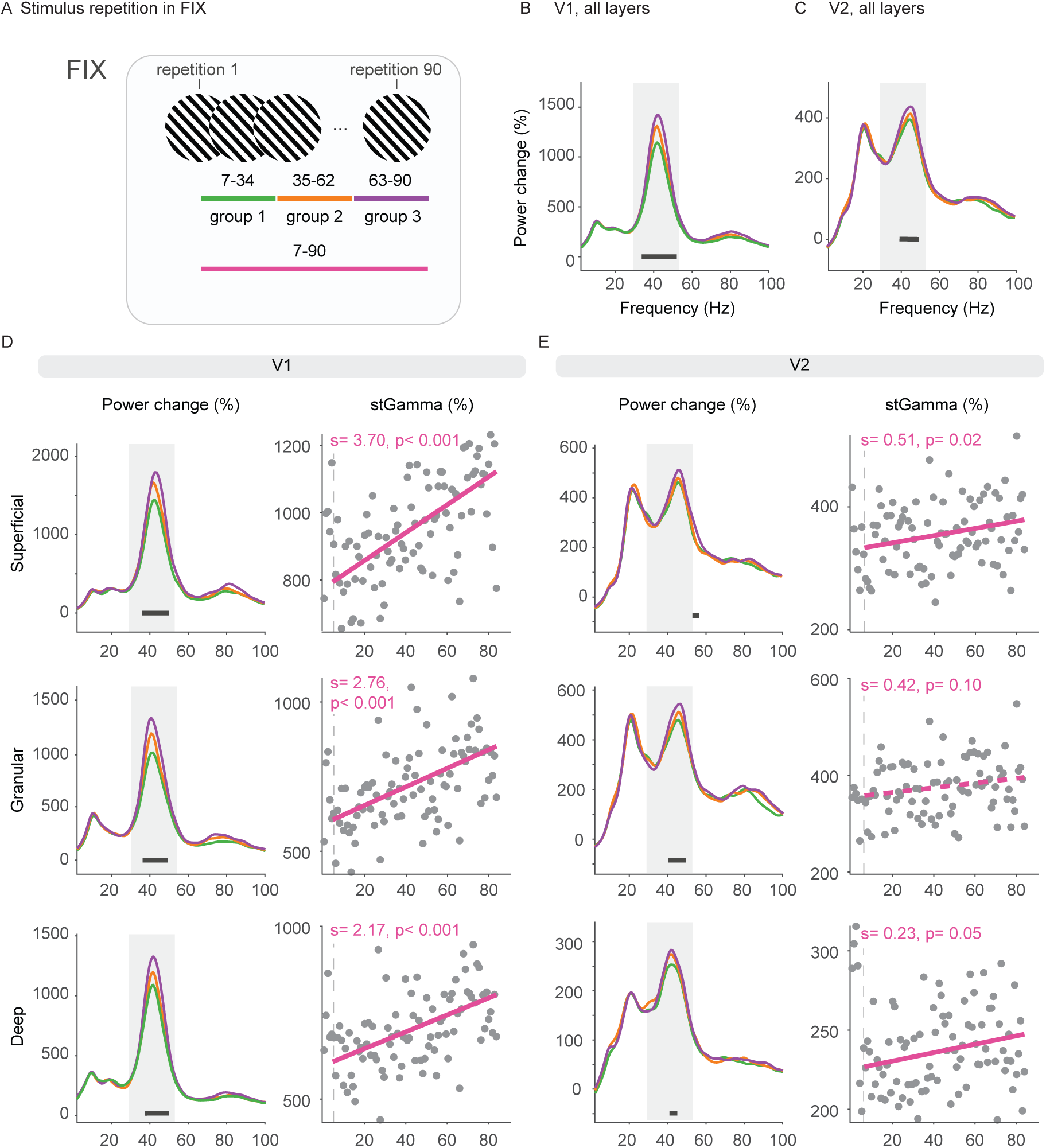
Repetition-related changes in bLFP power during FIX. (A) Schematic representation of the stimulus repetitions in FIX and their grouping into repetition groups 1, 2 and 3. Each group consisted of 28 repetitions. For the analysis of stGamma in D and E, we also defined a Trial-7-90 group. (B-C) Power change from baseline averaged over all V1 (B) and V2 sites (C), per repetition group (color-coded as in A). (D) Left: Power change from baseline per repetition group per laminar compartment (from top to bottom: superficial, granular, and deep). Grey rectangles indicate the gamma frequency range that was used to estimate the stGamma responses. Horizontal black bars indicate frequency ranges of significant difference between groups 1 and 3.Right: stGamma as a function of repetition number within FIX. The dashed grey vertical lines indicate the 7^th^ repetition. Regressions with (non-)significant slopes are plotted as (dashed) solid lines. The respective slopes (s) and p-values (p) are reported on top of each panel. (E) Same as in D but for V2. Note that y-axes were scaled per plot to optimally illustrate the effects.

### The repetition of a fixed-orientation grating stimulus leads to gamma-power increases and spike-rate decreases across layers

We investigated the repetition-related effects of a fixed-orientation grating stimulus (FIX block) on the amplitude of gamma-band power and the spike rates (MUA). To this end, we employed two complementary approaches.

First, to localize repetition effects on MUA in time, we compared time courses between an early and a late group of trials in the FIX block. Of the 90 trials of the FIX block, the initial 6 were excluded, and the remaining 84 were divided into three non-overlapping groups of 28 trials (Fig. 2A). We compared the first and third of those 28-trial groups to assess the repetition effect. The exclusion of the initial 6 trials was motivated by the previous finding that gamma-band power, analyzed below, often shows a distinct behavior in response to the first few repetitions of a given stimulus, with likely a different mechanism compared to the later repetition-related increase [4, 8; see Discussion]. Figure 2B and C illustrate the mean MUA responses for all V1 and V2 recording sites and for each trial group. Comparing trial group 3 versus group 1, MUA did not show significant differences in area V1 (Fig. 2B, D), whereas there were some significant differences during the early transient in area V2, both for the MUA pooled over all compartments (Fig.2 C) and for the superficial and granular compartments (Fig. 2E).

Second, we obtained single-trial estimates that allowed us to look for neuronal changes over the course of stimulus repetitions. Single-trial estimates were obtained by averaging MUA responses of a given trial over a time window (see Methods for a detailed description). We refer to single-trial estimates of MUA responses relative to baseline as stMUA (see Methods for details). stMUA estimates were obtained by averaging MUA responses from each trial within a time window of 0.05 to 0.1s relative to stimulus onset. Visual inspection of stMUA in both areas V1 and V2 revealed two distinct patterns: (1) a sharp decline during the initial approximately 20 trials, followed by (2) a period of more subtle changes throughout the subsequent 70 trials. To capture both patterns, we applied two separate linear regression models: one for the first 20 stMUA responses (referred to as “Early-20”) and one for the stMUA responses from trials 21-90 (referred to as “Remaining-70”), as illustrated in Fig. 2A. The regression slopes were significantly negative for all V1 and V2 laminar compartments during the Early-20 trials, and for the superficial compartments of V1 and V2 as well as the granular V2 during the Remaining-70 trials (Fig. 2D-E). When we considered the stMUA pooled over all sites in V1 and separately in V2, the regression slopes were significantly negative for the Early-20 in both areas and the Remaining-70 trials in V2 (Fig. S3).

We then performed corresponding tests for the repetition-related effects on LFP power. In accordance with previous studies [3, 4, 8], we observed a significant increase in gamma-band power from group 1 to group 3 in both areas V1 and V2 (Fig. 3B-C). This increase was not only evident in the grand averages of V1 and V2 but also within each laminar compartment (Fig. 3D-E).

Given that the repetition-related increase was confined to the gamma band (34-53 Hz for V1 and 38-49 Hz for V2; Fig. 3B,C), our subsequent single-trial analysis was focused on this frequency range. The single-trial estimates of LFP power in the gamma-frequency range are referred to as stGamma. Note that we defined the different frequency bands on the basis of the comparison between stimulus and baseline (averaged over all FIX trials), and not on the basis of any repetition effects (see Methods and Fig. S2 for details and resulting band definitions). The resulting frequency band for gamma was 29-53 Hz. For this gamma band, stGamma was computed, and the stGamma values of trials 7 to 90 were fitted with a linear regression (a separate regression analysis of the initial 6 trials was not performed due to the small amount of data). The regression slopes were significantly positive for all laminar compartments of V1 (Fig. 3D) and the superficial and deep compartments of V2 (Fig. 3E) and when all compartments were pooled per area (Fig. S3). The first six trials were excluded from the regression fits for reasons mentioned earlier.

### Repetition-related changes in inter-laminar interactions

We then investigated the effects of stimulus repetition on the information flow between laminar compartments inside V1 and inside V2. Information flow was quantified by computing Granger causality (GC) between all possible intra-areal inter-laminar site pairs and subsequent averaging for each combination of laminar compartments, separately for each direction of information flow. To test for repetition-related GC changes, we used the FIX block and compared trial group 1 to group 3. Similar to our analysis of power spectra, we first compared GC spectra per frequency. However, we noticed that the randomization distributions were dominated by GC values in the lower frequency range, which could mask putative effects in higher frequencies. Therefore, we additionally present a less fine grained comparison, after first averaging GC values within the distinct frequency bands as defined in the Methods and shown in Figure S2. The results of both statistical approaches are reported in Figure 4, and the results for the separate bands are discussed here. In area V1, in the alpha-frequency range, there was an increase in GC with stimulus repetition from deep to superficial, deep to granular, superficial to deep and superficial to granular compartments. In the beta range, we observed a significant increase from granular to superficial and superficial to deep compartments. In gamma, there was a significant increase from granular to superficial, from deep to superficial and deep to granular layers. In area V2, the effects were mostly confined to the gamma-frequency band, with an increase from deep to superficial compartments.

**Figure 4.**
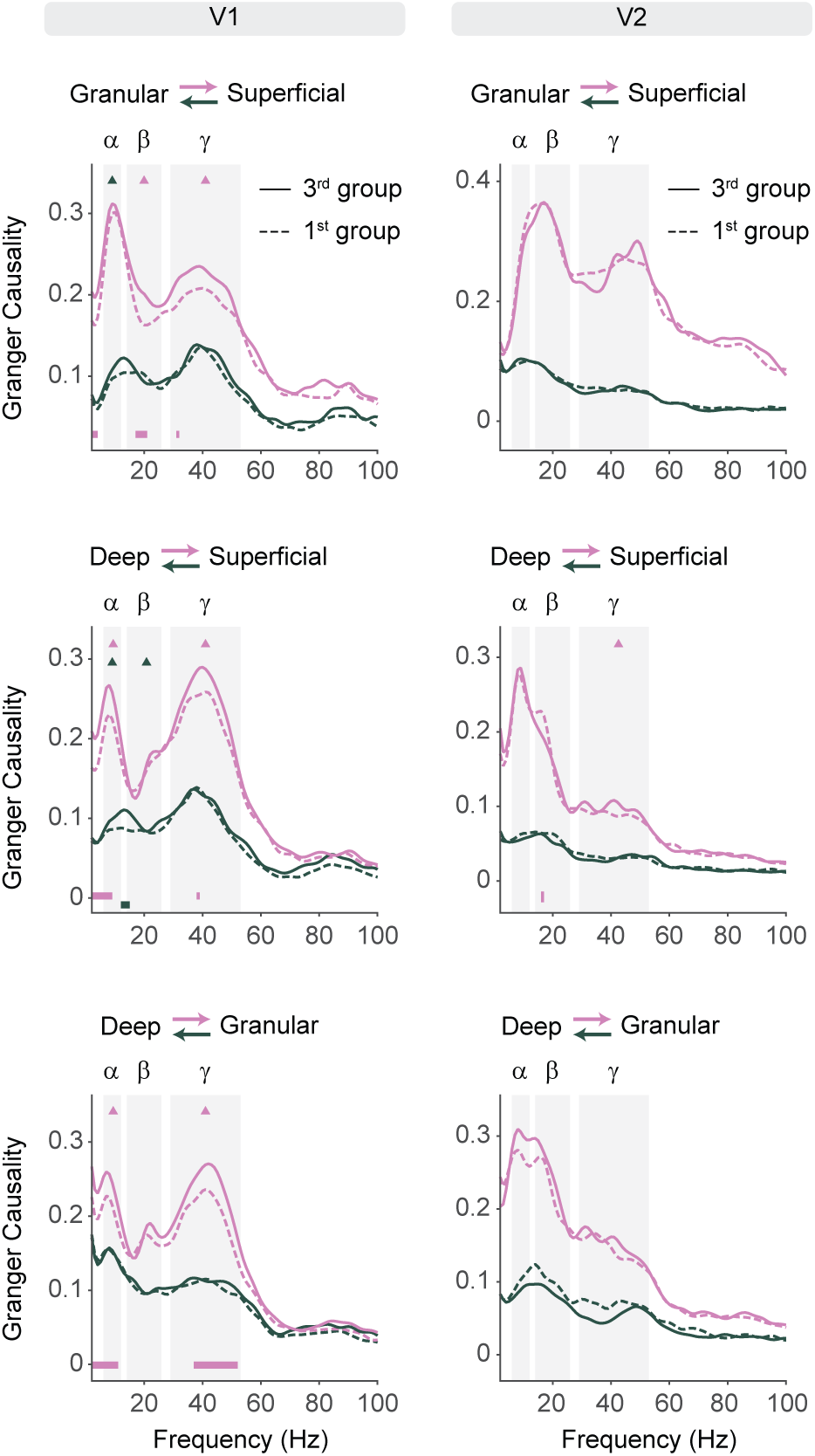
Repetition-related changes of Granger Causality between laminar compartments during FIX. Granger Causality (GC, see Methods for details) during the 1^st^ (dashed line) and 3^rd^ repetition group (solid line) of FIX. Different colors represent the GC direction as indicated on top of each panel. Grey rectangles show the three frequency bands that were used for the statistical analysis: alpha (α), beta (β), and gamma (γ). Triangles indicate statistically significant changes per frequency band, color-coded per GC direction; all significant changes were increases. Horizontal lines at the bottom of the plots (color-coded per directionality) indicate frequencies with significant differences derived from the statistical comparison across all frequencies.

Figure S4 shows a corresponding analysis for inter-areal GC between laminar compartments of V1 and V2. Note that the bLFP-RFs of the simultaneously recorded V1- and V2-sites typically overlapped only to a small degree or not at all, and therefore (1) gamma-band interactions were expected to be weak or absent, (2) the observed effects might be specific for interactions between V1 and V2 sites with non-overlapping RFs. Indeed, a clear gamma peak was mainly observed for GC from the deep V1 to the deep V2 compartment, maybe because deep-layer RFs are larger [35, 52]. Overall, feedforward GC from V1 to V2 often showed a clear alpha peak, whereas feedback GC from V2 to V1 often showed a clear beta peak. The repetition effects showed clear patterns that differed between alpha and beta. For alpha, all significant repetition effects were increases. By contrast, for beta, all significant repetition effects were decreases, and they all involved the granular or deep V1 compartments.

### The repetition of multiple interleaved stimuli leads to MUA and gamma-band changes

So far, we demonstrated that the repetition of a single orientation leads to plastic changes of neuronal activity across laminar compartments of V1 and V2. We next asked whether these effects were also present when many different stimuli were presented interleaved (Fig. 5A). As the above analyses found qualitatively similar repetition-related results across the laminar compartments and in both areas, all following analyses will pool the laminar compartments per area, and we add for most analyses the results for the pooled areas, referred to as V1+V2. We did this to simplify the presentation and enhance sensitivity.

**Figure 5.**
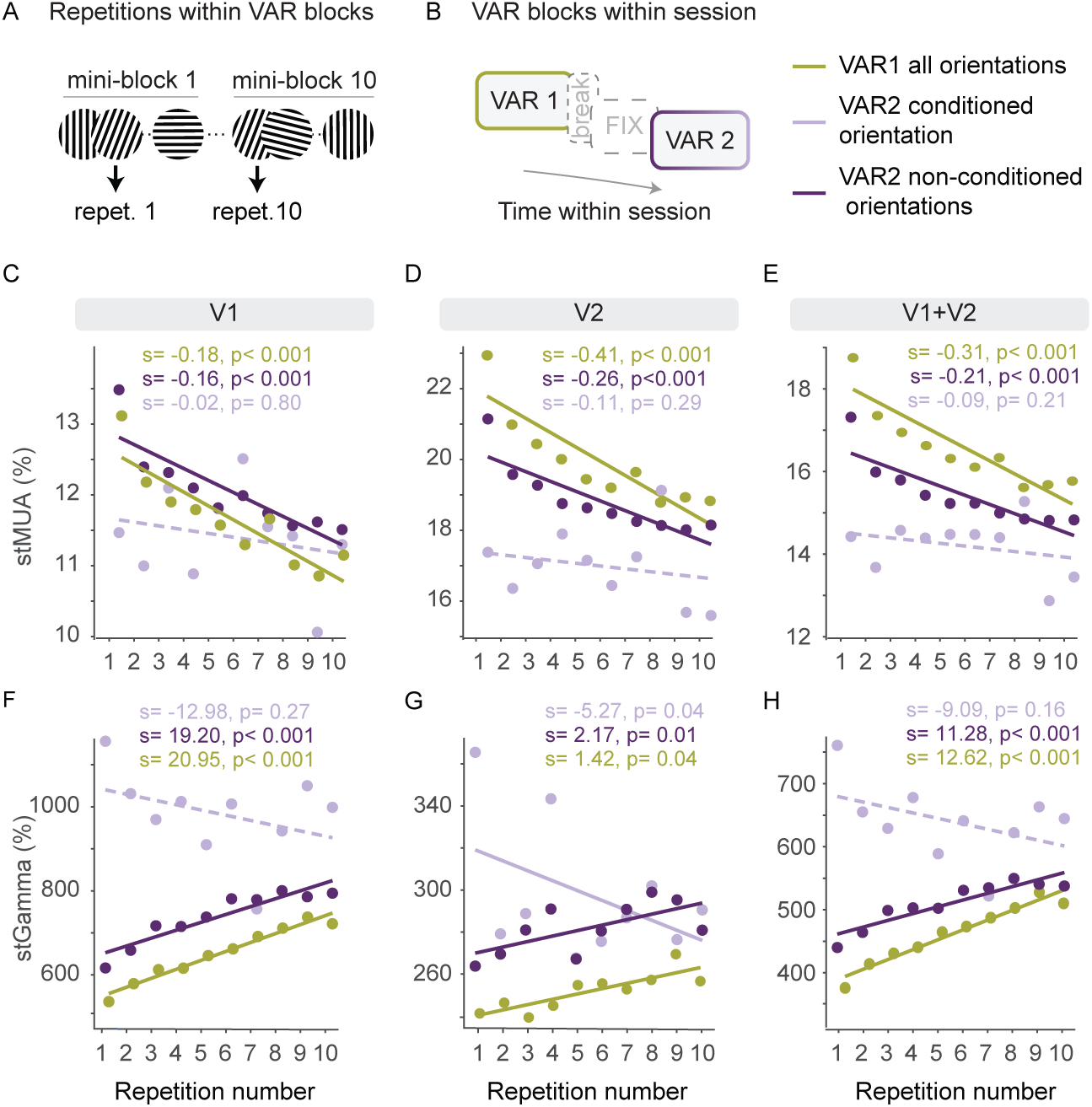
Repetition-related changes during VAR blocks in V1 and V2. (A) Schematic representation of stimulus repetitions across mini-blocks in VAR, illustrating that a given orientation occurred at a random position per mini-block. (B) Block sequence within a recording session. (C - E) stMUA as a function of the repetition number of a given stimulus orientation across the mini-blocks. For VAR1, neuronal responses were averaged over all orientations. For VAR2, neuronal responses were divided into two groups: responses for non-conditioned orientations and responses for the conditioned orientation. Each group is color-coded and presented for V1 (C), V2 (D) and combined V1+V2 (E). Lines indicate the respective linear fits (solid (dashed) for (non-)significant slopes). Slopes (s) and p-values (p) are color-coded and reported at the top of each panel. (F - H) Same as C-E, but for stGamma. Results of the statistical comparisons of repetition-related stMUA changes between conditioned and non-conditioned orientations in VAR2 (listed are the p-values): (1) difference between slopes: V1: 0.056, V2: 0.168, V1+V2: 0.096; (2) difference between intercepts: V1: < 0.001, V2: < 0.001, V1+V2: < 0.001. Results of the statistical comparisons of repetition-related stGamma changes between conditioned and non-conditioned orientations in VAR2 (listed are the p-values): (1) difference between slopes: V1: 0.01, V2 < 0.001, V1+V2: 0.004; (2) difference between intercepts: V1 < 0.001, V2 < 0.001, V1+V2 < 0.001.

We first investigated repetition-related plastic changes in VAR1. In this block, eighteen grating orientations were repeated interleaved. Importantly, it was the first block that the monkey completed in each session (Fig. 5B). This ensured that any observed repetition-related neuronal changes (within the session) could only result from repetitions within this block, excluding the influence from the FIX block. VAR1 consisted of ten mini-blocks. In each mini-block, all eighteen orientations were presented once in a randomized order. This design prevented immediate repetitions of the same orientation within a mini-block, while allowing each orientation to be repeated across the ten mini-blocks (Fig. 5A).

stMUA and stGamma responses were calculated per orientation and mini-block, resulting in ten single-trial estimates per orientation. For each orientation, the ten estimates were first fitted with a linear regression model. All estimates and fits were then averaged, leading to a grand-average fit.

Interestingly, stMUA and stGamma of V1, V2, and V1+V2 showed qualitatively similar repetition effects as observed in the FIX block. The slope of stMUA was significantly negative (green data in Fig. 5C-E), indicating a decrease in spiking activity with stimulus repetition. Conversely, the slope of the stGamma was significantly positive (green data in Fig. 5F-H), indicating an increase with repetition.

Thus, changes in MUA and gamma as observed with many repetitions of a single orientation in FIX, were also seen during VAR1 with 18 different orientations interleaved. Note, however, that the experimental design of VAR1 did not allow us to dissociate between the repetition of a specific orientation versus the overall repetition of stimuli across mini-blocks. Previous results suggest that when different orientations are presented in separate adjacent blocks, there is some overall gamma increase across the session, yet the dominant effect is a repetition-related stimulus-specific gamma increase [8]. In the following, we will use VAR2 to investigate whether such stimulus-specificity is also present for 18 interleaved orientations.

### The repetition of multiple interleaved stimuli leads to stimulus-specific neuronal changes

VAR2 had the same structure as VAR1, but it was completed immediately after FIX. This allowed us to investigate lasting effects of the orientation that had been conditioned during FIX. We looked for plastic changes in VAR2 separately for the conditioned orientation and for all other orientations combined, referred to as non-conditioned orientations. We hypothesized that different repetition-related response patterns between these two groups could indicate stimulus specificity.

Note that this analysis led to unequal numbers of trials per group. For the conditioned orientation, single-trial estimates were based on only one trial per mini-block. For the non-conditioned orientations, they were based on 17 trials. This probably led to higher noise levels in the estimates for the conditioned orientation.

In V1, V2 and V1+V2, the stMUA slopes for the non-conditioned orientations were significantly negative, whereas the stMUA slopes of the conditioned orientation did not show a significant effect (Fig. 5C-E). Direct comparison of the regression lines for the conditioned versus non-conditioned orientations revealed that the slopes were almost significantly different for V1 (p=0.056), but not for V2 or V1+V2. Intercepts were significantly lower for the conditioned orientation for V1, V2 and V1+V2 (see Fig. 5 legend for details of the statistical tests).

stGamma responses for the non-conditioned orientations showed the expected positive slope for V1, V2 and V1+V2 (Fig. 5F-H). By contrast and intriguingly, stGamma responses for the conditioned orientation showed a negative slope for V2, and a non-significant negative slope for V1and V1+V2. Direct comparison of the regression lines for the conditioned versus non-conditioned orientations revealed that, for V1, V2 and V1+V2, the conditioned orientation showed higher intercepts and lower slopes (see Fig. 5 legend for details of the statistical tests).

Thus, stGamma responses to the conditioned versus the non-conditioned orientation differed both at the beginning and during the first ten mini-blocks of VAR2. This demonstrates that repetition-related effects (induced during FIX) did not only survive the interleaving of 18 different orientations (during VAR2), but that they were at the same time stimulus specific, i.e., different between the conditioned and non-conditioned stimulus. Furthermore, there was partial persistence over time. The difference between stGamma for conditioned versus non-conditioned orientations decreased over the course of the ten mini-blocks, such that by the 10^th^ mini-block, the two orientation groups elicited responses of similar amplitude (Fig. 5F-H). This implies a partial persistence of the gamma-band increase induced during FIX despite the interleaved repetition of 17 other stimuli during VAR2. Taking into account the trial duration of VAR2, the time period to complete all ten mini-blocks ranges between at least 8.7 and 9.3 minutes, without considering additional delays due to incorrect trials like fixation beaks.

### Gamma-band plasticity is highly stimulus specific

We next quantified the degree of stimulus specificity of the repetition-related changes induced by the FIX block. Do these repetition effects transfer to orientations that are close to the conditioned one? By comparing the neuronal responses in VAR2, that is after FIX, to the responses in VAR1, that is before FIX, we were able to investigate the effects of conditioning one orientation on the neuronal responses for all orientations, including the conditioned orientation itself, immediately neighboring orientations and more dissimilar orientations.

On each recording day, the conditioned orientation was chosen randomly, leading to different conditioned orientations across sessions. To combine data across sessions, we expressed all orientations as a function of their orientation difference from the conditioned orientation. As we had no reason to believe that positive and negative orientation differences had different effects, we pooled corresponding absolute orientation differences, leading to orientation differences ranging between 0 and 90 °.

As we showed earlier, there are plastic changes that take place during VAR blocks due to repetitions across mini-blocks. We aimed at isolating the effect of the conditioning happening in FIX while eliminating the influence of VAR effects. To this end, we estimated the responses at the end of VAR1 and at the start of VAR2, each time using all data of the respective blocks, but eliminating the changes that occurred within those blocks. Specifically, for each orientation, we fitted linear regressions to VAR1 responses and used the value of this regression at the final VAR1 stimulus presentation as the best estimate of the last response just before FIX, referred to as Pre-conditioning value, or just “Pre” in Fig. 6. Correspondingly, we fitted linear regressions to VAR2 and used the intercept of this regression, i.e. its value for a hypothetical zeroth presentation before any effect of the first VAR2 mini-block, referred to as Post-conditioning value or just “Post” in Fig. 6. Then, we computed the difference of Post minus Pre as an estimate of the effects induced by the intervening FIX.

**Figure 6.**
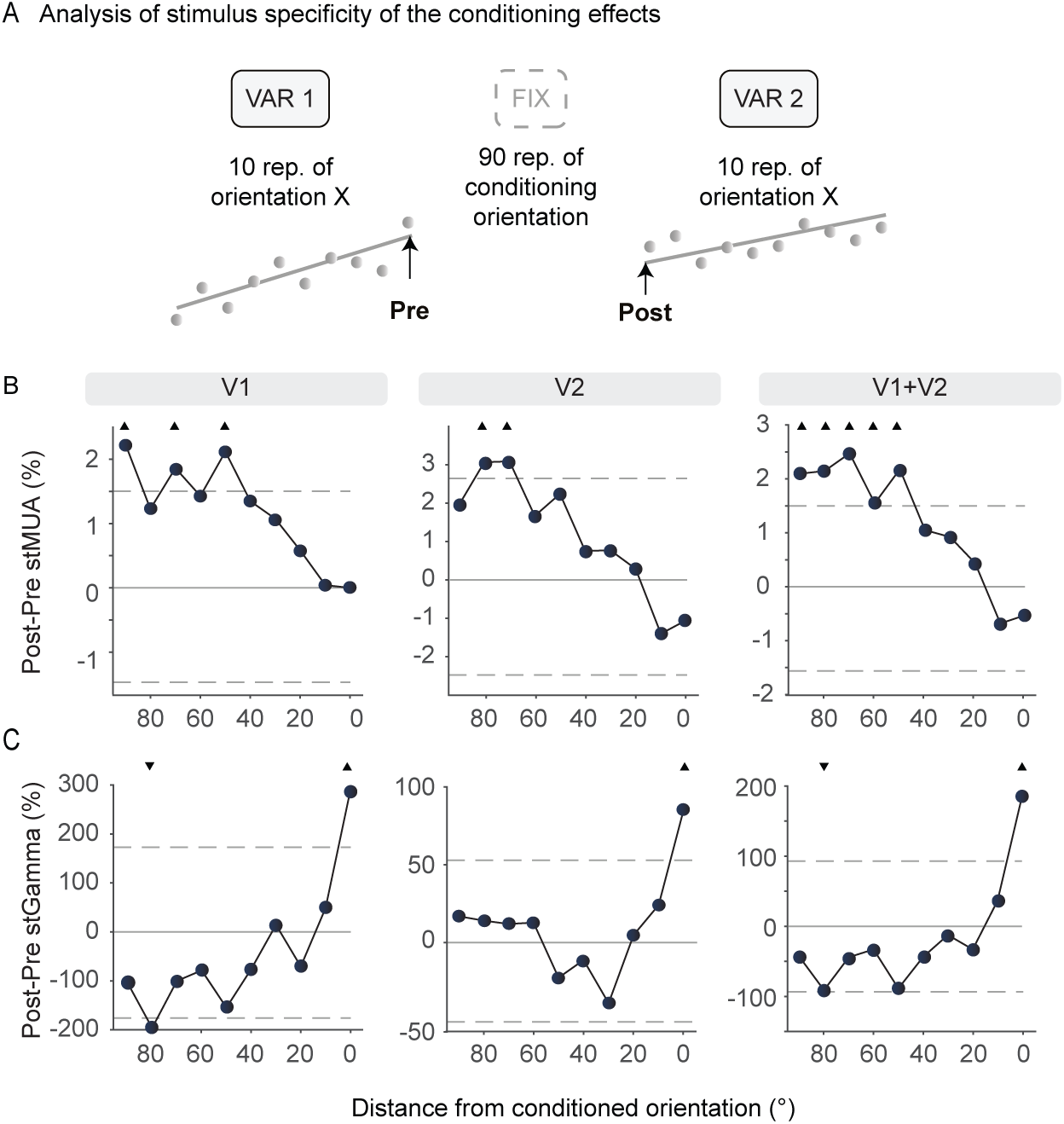
Stimulus specificity of the repetition-related changes. (A) Schematic representation of the analysis of stimulus specificity. For each orientation, single-trial estimates of the neuronal responses were fitted with linear regressions in VAR1 (pre conditioning) and VAR2 (post conditioning). The conditioning-related change of the neuronal responses was then calculated as the difference between the intercept of the VAR2 fit (marked as “Post”) minus the last point of the VAR1 fit (marked as “Pre”). The conditioning-related change (Post-Pre) for stMUA (B) and stGamma (C) are plotted as a function of orientation difference from the conditioned orientation. Positive and negative orientation differences have been combined. Dashed horizontal lines indicate the upper and lower significance limits derived from the randomization distribution. Triangles indicate orientation differences with statistically significant conditioning effects, with the triangles pointing upward for increases and downward for decreases.

The conditioning-induced stMUA changes were qualitatively similar in V1 and V2 (Fig. 6B). There seemed to be a tendency for an increase with orientation distance from the conditioned orientation. This increase reached significance for several of the orientations with large distance to the conditioned orientation.

The conditioning-induced stGamma changes were again qualitatively similar in V1 and V2 (Fig. 6C), but dissimilar from the stMUA changes. For gamma, there was a significant increase at the conditioned orientation, for V1, V2 and V1+V2, and no significant changes for all but one of the other conditions in V1 and in V1+V2. Overall, the pattern was consistent with an increase that is specific for precisely the conditioned orientation.

Note that VAR1 and VAR2 were separated by substantial time, for the break between VAR1 and FIX, and for FIX itself. With this passage of time, we expected some decay of the VAR1-induced changes between Pre and Post (except of course for the conditioned orientation; Peter, Stauch (4)). Given that VAR1 induced MUA decreases and gamma increases, the decay of these effects should lead to opposite effects on the Post-minus-Pre difference (i.e. our estimate of the conditioning during FIX): The MUA effects should be slightly shifted upwards, whereas the gamma effects should be slightly shifted downwards. Indeed, for MUA, most significant effects were increases, and for gamma in V1, there was one unexpected significant decrease.

As a control, we repeated the same analyses, yet not aligned to the conditioned orientation but to the preferred orientation of the respective recording site. As mentioned above, the conditioned orientation was randomly chosen per session to eliminate confounds with the preferred orientation. The analyses aligned to the preferred orientation revealed several significant changes for stMUA (Fig. S5B). For stGamma, there were no significant changes, except for one orientation in V1 (Fig. S5C). These significant results were increases for stMUA and one decrease for stGamma. Thereby, they are consistent with the abovementioned decay for the non-conditioned orientations due to time passing during FIX. Overall, these control analyses suggest that the experimental design was successful at avoiding a confound between the conditioned and the preferred orientation.

### Neuronal responses across trial blocks suggest effects of temporal stimulus predictability

When we compared the spectral response to the conditioned orientation during FIX and VAR2, we found that the strength of the alpha and beta rhythms was conspicuously higher during FIX than VAR2 in both areas and all laminar compartments (Fig. 7B and Fig. S6). When we investigated single-trial responses across all trials, including all orientations, of a session, we noticed intriguing dynamics across trial blocks (Fig. 7C-E). This was particularly conspicuous for alpha and beta power, and since these two frequency bands showed similar effects here, we pooled them in a combined alpha-beta band (6-26 Hz) for the remaining analyses and refer to these single-trial estimates as stAlpha-Beta. In area V1, stAlpha-Beta showed in each block, for approximately the first 20 trials, a strong increase, and for the remaining trials of the block it either stayed at this high level or decreased slightly (Fig. 7C, see Fig. S7 for similar results in area V2). Note that for this and the following analyses, to show the stability of the effects, we included all trials that the monkey completed in VAR2 up to the minimal number available in all sessions. For FIX, alpha-beta power rose to and stayed at a substantially higher level than for VAR1 or VAR2. Table 2 reports the p-values of the statistical comparisons between the means of the regressions of each block. Note that in FIX, the conditioned orientation was repeated in each trial, rendering it highly predictable, whereas in VAR, the 18 different orientations were randomly interleaved, rendering a given orientation much less predictable. Studying this orientation predictability was not the purpose of our experimental design, such that the design was not optimized accordingly, but nevertheless, the results seemed intriguing and potentially inspiring for more targeted future investigations.

**Figure 7.**
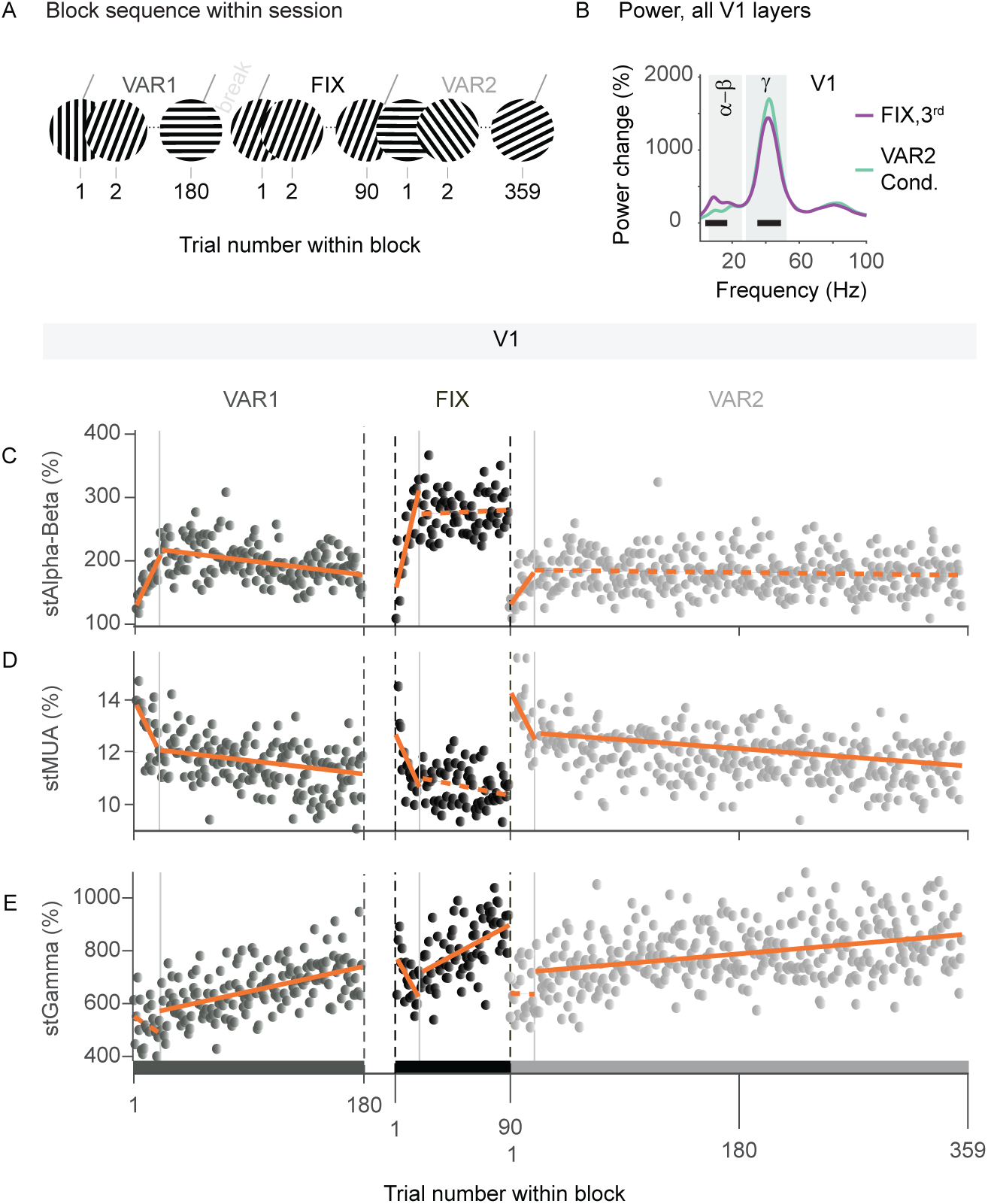
Comparison of neuronal responses in V1 across blocks. (A) Schematic representation of the block sequence within the session, and the trial number within the block. Note that for this and the following analyses, we included all trials that the monkey completed in VAR2 up to the minimal number available in all sessions. (B) Power change relative to baseline in response to the conditioned orientation in the 3^rd^ repetition group in FIX (purple) and in VAR2 (light green). Black horizontal lines show frequencies with significant difference between the conditions. Single-trial estimates for alpha-beta (C), MUA (D), and gamma (E) are plotted for all trials (irrespective of stimulus orientation) as a function of trial number. Orange lines show the two regressions fitted per block for trials 1-20 and trials 21 until the end of that block. Solid (dashed) lines indicate that the slopes of the respective linear fits are (not) significantly different from zero. Grey vertical lines show the 20^th^ trial in each block. The slopes and p-values are summarized in Table S1. A statistical comparison of the regression means are presented in Table 2.

**Table 2.**
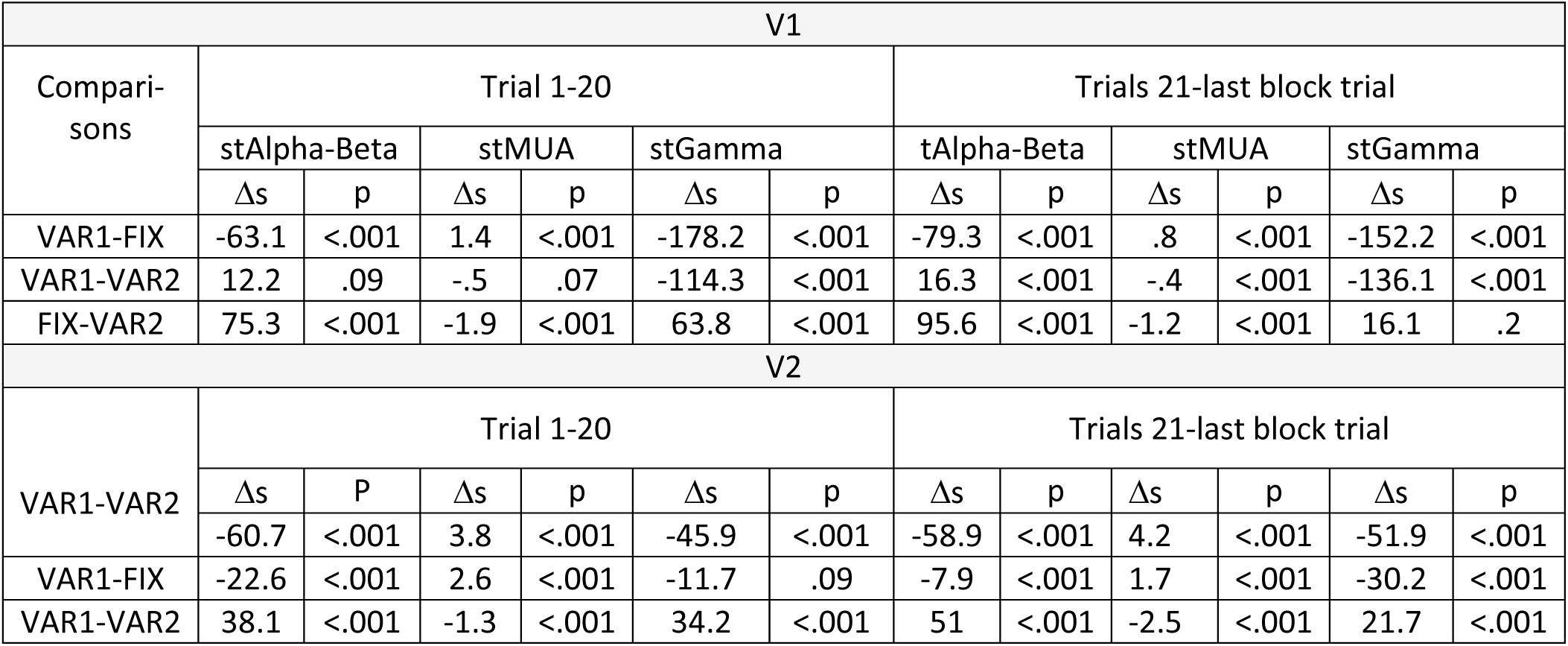
Statistical comparison of the mean values of the regression fits between blocks. Reported are difference of the regression means (Δs) and the corresponding p-values (p).

| V1 |  |  |  |  |  |  |  |  |  |  |  |  |
| --- | --- | --- | --- | --- | --- | --- | --- | --- | --- | --- | --- | --- |
| Compari-<br>sons | Trial 1-20 |  |  |  |  |  | Trials 21-last block trial |  |  |  |  |  |
|  | stAlpha-Beta |  | stMUA |  | stGamma |  | tAlpha-Beta |  | stMUA |  | stGamma |  |
| | $\Delta s$ | p | $\Delta s$ | p | $\Delta s$ | p | $\Delta s$ | p | $\Delta s$ | p | $\Delta s$ | p |
| VAR1-FIX | -63.1 | <.001 | 1.4 | <.001 | -178.2 | <.001 | -79.3 | <.001 | .8 | <.001 | -152.2 | <.001 |
| VAR1-VAR2 | 12.2 | .09 | -.5 | .07 | -114.3 | <.001 | 16.3 | <.001 | -.4 | <.001 | -136.1 | <.001 |
| FIX-VAR2 | 75.3 | <.001 | -1.9 | <.001 | 63.8 | <.001 | 95.6 | <.001 | -1.2 | <.001 | 16.1 | .2 |
| V2 |  |  |  |  |  |  |  |  |  |  |  |  |
| VAR1-VAR2 | Trial 1-20 |  |  |  |  |  | Trials 21-last block trial |  |  |  |  |  |
| | $\Delta s$ | P | $\Delta s$ | p | $\Delta s$ | p | $\Delta s$ | p | $\Delta s$ | p | $\Delta s$ | p |
|  | -60.7 | <.001 | 3.8 | <.001 | -45.9 | <.001 | -58.9 | <.001 | 4.2 | <.001 | -51.9 | <.001 |
| VAR1-FIX | -22.6 | <.001 | 2.6 | <.001 | -11.7 | .09 | -7.9 | <.001 | 1.7 | <.001 | -30.2 | <.001 |
| VAR1-VAR2 | 38.1 | <.001 | -1.3 | <.001 | 34.2 | <.001 | 51 | <.001 | -2.5 | <.001 | 21.7 | <.001 |

stMUA showed in all blocks, for approximately the first 20 trials, a strong decrease; this reached significance in all blocks (Fig. 7D). stMUA showed further decreases for the remaining trials of the VAR1 and VAR2 blocks, and the same trend for the FIX block. For FIX, stMUA decreased to and stayed at lower levels than for VAR1 or VAR2 (Table 2). This is again suggestive of an effect of stimulus predictability.

As mentioned above, previous studies had described that also stGamma can show decreases for the initial few repetitions of a given stimulus. The reports so far have described this decrease to occur across the initial 4 [4] or initial 10 [8] trials, and primarily for stimuli that were less overtrained than the gratings for the monkey used here. Nevertheless, to be consistent with the analyses of stApha-Beta and stMUA, we also analyzed stGamma separately for the initial 20 trials of each block and the remaining trials of each block (Table 2). For most blocks, stGamma across the initial 20 trials did not show a significant change; only for FIX in V1, a significantly negative slope was observed. By contrast, for the remaining trials, stGamma showed significant increases within each block and for both areas. The means of stGamma were higher in VAR2 than VAR1, and highest in FIX.

## Discussion

### Summary

- In accordance with previous studies, we found stimulus-specific MUA decreases and gamma increases with stimulus repetition [4, 8].
- We systematically investigated the level of stimulus specificity by presenting 18 orientations spaced by 10 ° before and after conditioning. By comparing neuronal responses before and after conditioning, we found that the conditioning effect in gamma is highly specific for the conditioned orientation.
- The repetition-related changes in gamma and MUA existed in all recorded laminar compartments.
- Repetition-related changes in gamma and MUA were evident even when 18 different orientations were interleaved. This suggests that these effects might generalize to natural viewing conditions, where also many stimuli are expected to be interleaved.
- At the same time, we found clear indications that these changes were still stimulus specific, as VAR2 showed clear differences in intercept and slope of the regression for gamma between conditioned and non-conditioned orientations.
- Finally, we report changes of alpha-beta power and MUA across blocks that suggest a potential relation to stimulus predictability.

### Limitations

The data presented here is from one monkey, which limits the inference to this sample. Note that almost all studies with awake macaques use few monkeys, i.e. typically two or three, and therefore, their inference is also limited to their sample, rather than to the population from which the sample has been drawn. Thus, because the inference remains qualitatively the same for studies with three, two or one animal, we have argued on grounds of ethical considerations that the publication of monkey studies should go forward also with one animal [48].

Furthermore, our test of stimulus specificity was performed with a resolution of 10 °, as we presented orientations spaced by this amount. Future studies might test finer steps, at least close to the conditioned orientation. We had opted against an uneven sampling of orientations during VAR1 and/or VAR2, because this would have also changed the probabilities of the different orientations, with unknown consequences for neuronal responses. The alternative, an evenly spaced sampling with smaller orientation steps would have resulted in a very large number of conditions and thereby a reduction in our sensitivity to observe effects. Yet, having established the effect here, future studies might be able to investigate the effect further, at higher resolution.

### Gamma changes on different timescales

Previous studies of repetition-related gamma changes have reported (1) a strong gamma response for the first presentation of a given stimulus that rapidly declines over the course of the subsequent few (up to 10) repetitions [4, 8, 53, 54], (2) a later, slower gamma increase that builds up over the course of further repetitions [3, 4, 8]. These two patterns of gamma changes might reflect the superposition of two distinct dynamics, corresponding to two processes [4, 8]. Specifically, it has been suggested that the strong initial gamma response reflects novelty detection, and the late gamma increase is related to synaptic plasticity [4, 8].

In the present study, we intended to investigate the late gamma increase. Therefore, in the analysis of FIX, we excluded the first 6 trials. Regarding the VAR blocks, each mini-block contained 18 different orientations, and thereby a putative novelty-related response might not show the same degree of decrease as seen in previous studies with repetitions of identical or very few distinct stimuli [4, 8]. Our analysis of VAR based on the repetitions of a specific stimulus across mini-blocks indeed showed no indication of a novelty effect for the first mini-block. In any case, in the present data, any novelty effect in gamma seemed weak, even in FIX, probably because the animals were highly overtrained on the very grating stimuli used during data collection.

### Repetition-related gamma increases occur in all laminar compartments of V1 and V2

While some studies indicate that visually-driven gamma synchronization is more prominent in superficial cortical layers [20, 21, 55], others report additional peaks in deep and granular layers with a relative absence in the V1 input layer L4C [22, 23, 25, 29, 56]. We found prominent visually induced gamma-band power in the bLFP from all laminar compartments of both V1 and V2. Note that the V1 granular compartment combined L4C with other L4 sublayers, such that the gamma observed for the entire granular compartment might primarily originate from outside L4C.

We investigated for the first time the laminar distribution of the late repetition-related increase of gamma. Interestingly, we found prominent repetition-related gamma increases in all laminar compartments of V1 and V2.

### Potential mechanisms of late gamma-band increase

Stauch, Peter (8) suggested that the late repetition-related gamma increase might reflect plastic changes of the synaptic connectivity between excitatory and inhibitory neurons, and we will review their arguments here. Changes in the synaptic weights between excitatory and inhibitory neurons could emerge through the interaction between Hebbian spike-timing-dependent plasticity (STDP) and the spike timing in the gamma cycle that depends on the neuron’s stimulus drivenness. Neurons in early visual cortex are tuned to specific features like orientation. For a given stimulus orientation, the most strongly stimulus-driven neurons tend to spike earlier in the gamma cycle than the most weakly driven ones [57, 58]. Furthermore, within the gamma cycle, excitation is followed by inhibition within few milliseconds [59–62]. Together, these empirical results suggest that gamma-band activity induced by a particular stimulus entails an activation sequence of strongly stimulus driven excitatory neurons (E_strong_), followed by local inhibition (I_local_), followed by weakly stimulus driven excitatory neurons (E_weak_), or in short an E_strong_-I_local_- E_weak_ sequence [8]. During stimulus repetition, the same E_strong_-I_local_-E_weak_ sequence is repeatedly activated, and typically followed by reward, providing ideal conditions for Hebbian STDP [63, 64].

Importantly, synaptic weights between connected neurons can be either potentiated or weakened depending on the temporal relation of their spiking: Inputs from the leading to the lagging neurons are typically strengthened, whereas inputs from the lagging to the leading neurons are typically weakened [64]. Thereby, the repeated activation of E_strong_-I_local_-E_weak_ could lead to a strengthening of the connections from E_strong_ to I_local_ and from I_local_ to E_weak_. Strengthening of the connection from E_strong_ to I_local_ would allow the E_strong_ neurons to drive I_local_ more efficiently and faster, which could explain both the increase in gamma-band power and in the peak frequency [3, 4, 8]. A strengthening of the connections from I_local_ to E_weak_ neurons would lead to increased inhibition of the E_weak_ population, and thereby a sharpening of the neuronal population response. See Figure 6 in Stauch, Peter (8) for a schematic representation of the proposed mechanism.

The abovementioned mechanism [8] could explain the high stimulus specificity that we report in this study. Note that this mechanism does not place strong constraints on the spatial scale, as long as the involved neurons engage in gamma-band synchronization, which has been found locally [23, 56, 65–69] and interareally [56, 70–72]. Thus, these plastic changes could take place within a layer of an activated cortical column, across an entire column, and between connected columns within an area or even between areas.

Among those options, a particularly intriguing one is that the late repetition-related gamma increases could be due to plastic changes of the intrinsic, i.e. intra-areal, horizontal connections. Those horizontal connections originate from excitatory neurons [73], target both excitatory and inhibitory neurons [74], and they are orientation specific in superficial layers of V1 [75, 76]. Plastic changes of these connections underlie cortical reorganization in macaque and cat V1 following retinal lesions [77–79], and also perceptual learning (for a review see Gilbert and Li (80)). Importantly, horizontal connections have been suggested to play an important role in the generation of gamma oscillations [66, 67, 81, 82]. Electrophysiological recordings in cat V1 revealed that distant cortical columns with non-overlapping RFs synchronize in gamma only when they exhibit similar orientation preference [83]. The orientation-selective anatomical and functional connectivity predict that during the repetition of a specific orientation, cortical columns preferentially tuned to the presented orientation are repeatedly activated and synchronize in gamma, entailing spiking with precise relative timing. This might lead to a strengthening of horizontal connectivity with maintained orientation specificity.

In fact, a putative involvement of intrinsic horizontal connections might explain our observation that the repetition-related changes showed more pronounced orientation selectivity for gamma than for MUA responses. Gamma might be specifically dependent on long-range horizontal connections, which are particularly orientation selective. This orientation selectivity might thereby be transferred to gamma and its changes with stimulus repetition.

In line with the potential interaction between Hebbian plasticity and gamma oscillations, Galuske, Munk (84) found that the repeated presentation of a grating leads to plastic changes of orientation domains in primary visual cortex of anesthetized cats. Importantly, the induced rearrangements of the orientation domains were only present under conditions of strong visually induced gamma. Thus, gamma oscillations seem to play an important role in cortical plasticity.

As an alternative explanation for the observed repetition-related changes, one might consider plastic changes of thalamocortical connections. However, developmental studies have shown that thalamocortical projections do not show experience-dependent changes after the critical period [85]. Furthermore, gamma oscillations are cortically generated in awake non-human primates [86] and cats [87]. Therefore, plastic changes of the thalamic input are unlikely to explain repetition-related gamma changes.

Stimulus repetition could also lead to plastic changes of cortico-cortical feedforward influences, leading to more efficient feedforward signaling. A human MEG study found that stimulus repetition leads to strengthening of feedforward influences in gamma from lower to higher cortical areas both in the ventral and dorsal visual streams [8]. Such increases could be mediated by local changes that increase the postsynaptic impact of bottom-up signals, by increased feedforward connectivity or both.

Interestingly, a recent study from our lab used Dynamic Causal Modeling to investigate repetition-related effects in gamma activity in macaque areas V1 and V4. The best-fitting models included local changes within each area, as well as repetition-related changes in the gain of the neuronal inputs rather than the outputs [88].

Finally, top-down signals could also contribute to repetition-related changes in lower areas. Area V1 receives the majority of its feedback projections from V2 [89, 90], and interestingly, recent studies point to a crucial role of these V2-to-V1 projections in the generation of visually driven gamma and surround suppression in area V1. Reversible cooling of V2 drastically reduced visually driven gamma in V1 [91], and optogenetic inactivation of V2-to-V1 projections led to reductions in the surround suppression in V1 [92]. Note that considerations mentioned here for V1 and its interactions with higher areas can be similarly made for V2 and its interactions with areas above it.

### Effects of stimulus repetition and predictability in the context of predictive coding

Theories of predictive coding propose that the brain uses prior knowledge to predict upcoming events and sensory input. Sensory inputs are compared and subtracted from sensory predictions, resulting in a prediction error signal. This is thought to happen in a cascaded fashion across the levels of the visual hierarchy, with prediction-error signals being fed forward in the bottom-up direction, and predictions being fed back in the top-down direction through the respective anatomical projections [13–15].

This predictive-coding framework proposes that stimulus repetition leads to ‘representational sharpening’: Repeated stimuli become more predictable, and the more accurate predictions lead to reduced (precision-weighted) prediction errors, which are reflected in reduced firing rates. “Crucially, top-down predictions are not just about the content of lower-level representations but also about our confidence in those representations. This confidence may be mediated by modulating the postsynaptic gain of superficial pyramidal cells encoding prediction error—to boost their influence on higher levels. Mathematically, this gain corresponds to the precision (inverse variance) of prediction errors and provides a nice metaphor for attention (Feldman & Friston, 2010)” [quoted from 93]. In our data, the precision of prediction errors is enhanced by the conditioning protocol specifically for the conditioned stimulus. Furthermore, the postsynaptic gain of a neuronal group has been related to the gamma-band synchronization among the respective neurons [94] and the degree to which it entrains higher-area neurons [72, 95]. Taken together, this leads to the predictive-coding based hypothesis that gamma-band synchronization should be enhanced by the conditioning protocol specifically for the conditioned stimulus.

More generally, bottom-up and top-down interactions have been linked with distinct neuronal rhythms [26–30]. Several studies suggest that top-down predictions are mediated by alpha and beta oscillations [21, 96, 97]. Bastos, Lundqvist (21) found higher alpha and beta power for predictable compared to unpredictable stimuli, and Chao, Takaura (97) reported a beta-band power decrease during periods that required the update of sensory predictions. In line with these studies, we show that as a stimulus is repeated, alpha-beta power increases within a few trials and stays elevated until a prediction update is required, i.e. until an unexpected stimulus is presented (see transition from FIX to VAR2 in Fig. 7).

Gamma-band activity has been implicated in the bottom-up signaling of prediction errors, with stronger prediction errors leading to stronger gamma power [21, 97]. This effect has also been supported by previous stimulus-repetition studies [4, 8]: The first presentation of a novel stimulus led to a strong gamma-band response, which rapidly decreased over the course of the next few repetitions when the stimulus turned more predictable. Interestingly, these changes in gamma during the first few repetitions of a stimulus were prominent for novel [4, 8] but not for familiar stimuli like grating stimuli on which animals had been overtrained [4].

In the present study, we exclusively used overtrained gratings and confirmed this previous finding. We focused our investigation of gamma on the late repetition-related increase. This effect might also be explained in the context of predictive coding (see also Discussion in Katsanevaki, Bosman (88)). A generalization of the predictive coding framework posits that sensory inputs and their associated prediction errors are modulated by their inverse variance or precision [98, 99]. The precision of a prediction error is thought to be context dependent and to modulate the postsynaptic gain of the respective neurons, with increased precision leading to increased postsynaptic gain. The late repetition-related gamma-band increase could mediate this increased postsynaptic gain as precision rises with repetition. While spiking activity is decreased during repetitions, an increase in gamma synchrony could maintain or even boost the postsynaptic impact of the remaining spiking by means of increased feedforward coincidence detection [11] or by increased postsynaptic entrainment [100].

## Supplementary figure legends

**Figure S1.**
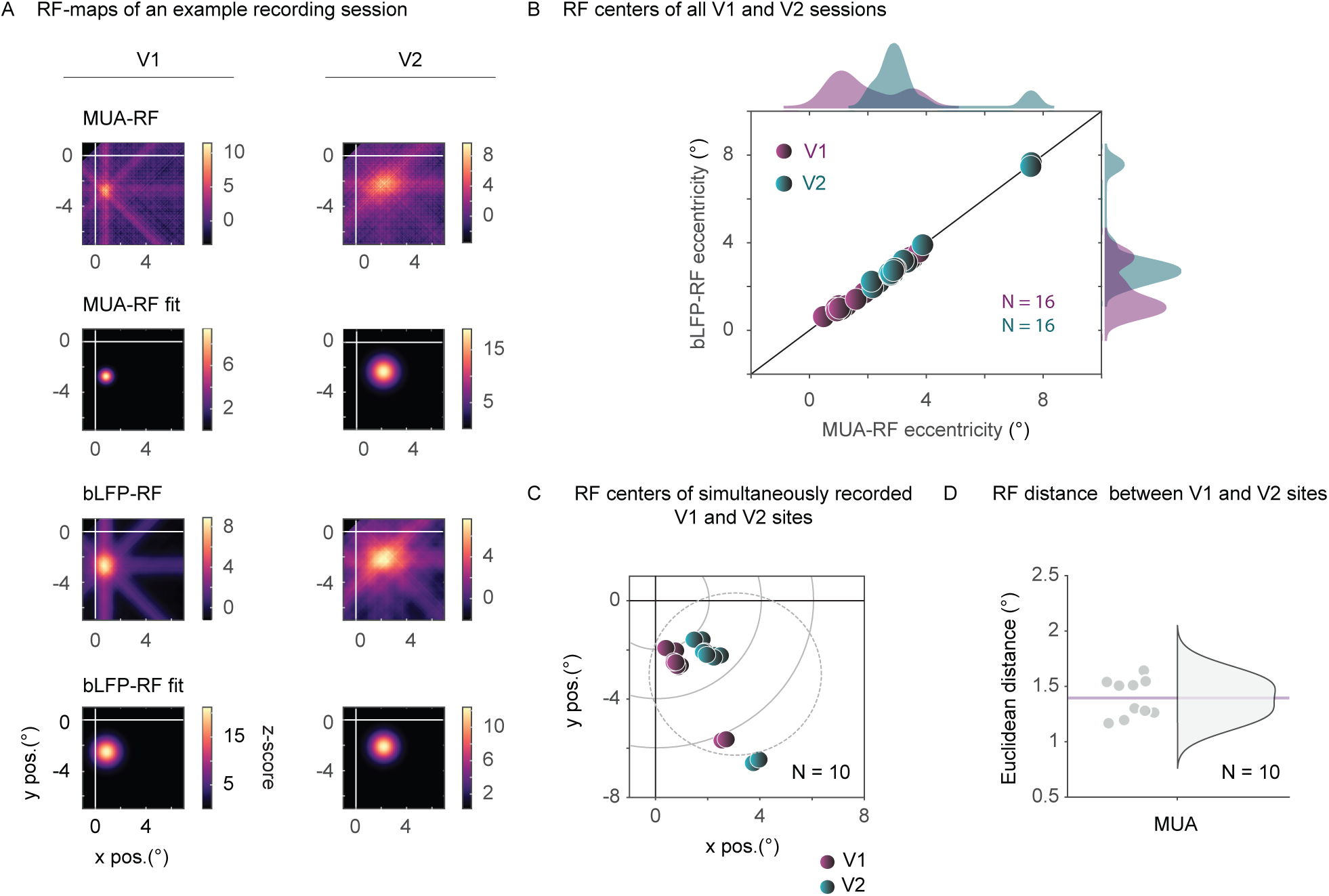
Examples and distributions of receptive fields (RFs). (A) MUA- and bLFP-RF maps of example recording sites in V1 (left) and V2 (right) and their respective Gaussian fits that were used to calculate the coordinates of the RF centers. For B-C, each dot represents the average over all contacts within one area in one session. (B) bLFP- and MUA-RF centers. Recording sessions in V1 (N=16) and V2 (N= 16) are color-coded in magenta and dark green, respectively. Note the strong similarity between bLFP- and MUA-RF centers. (C) MUA-RF centers of all sessions with simultaneous recordings in V1 and V2 (N=10). Same color-coding as in (B). (D) Euclidean distances between simultaneously recorded V1- and V2-RF centers shown in (C). The pink horizontal line indicates the mean Euclidean distance.

**Figure S2.**
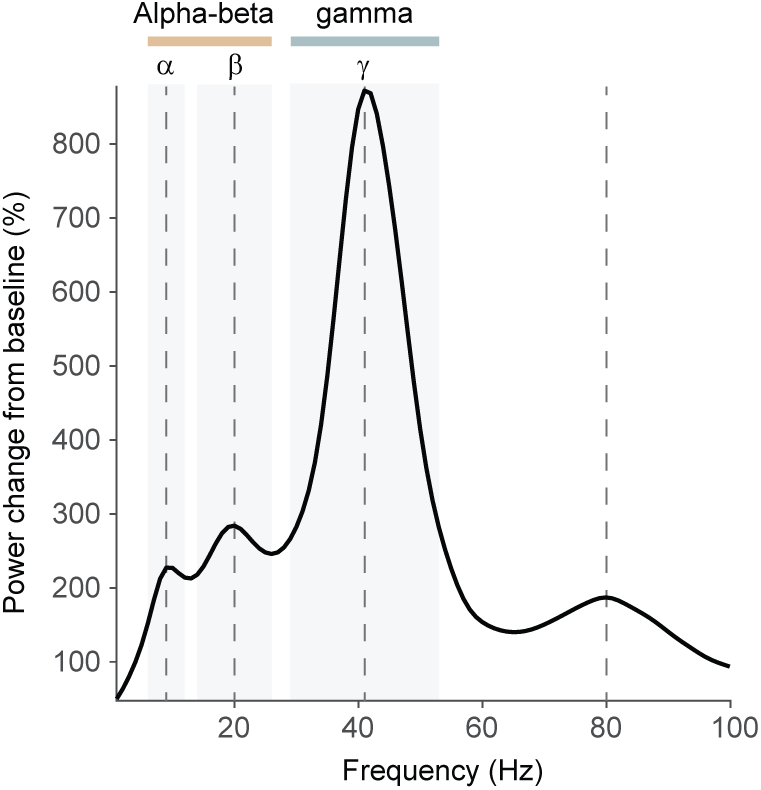
Definition of frequency bands. Power change relative to baseline averaged over all V1 and V2 recording sites. Dashed lines indicate the four distinct spectral peaks (at 9, 20, 41 and 80 Hz) that guided the definition of frequency bands. Based on these peaks, an alpha (6-12 Hz), a beta (14-26 Hz) and a gamma band (29-53 Hz) were defined (see Methods for details). Grey rectangles indicate each band and outline their respective frequency limits. Some analyses used a combined alpha-beta band (6-26 Hz), shown as a pink rectangle on top of the panel.

**Figure S3.**
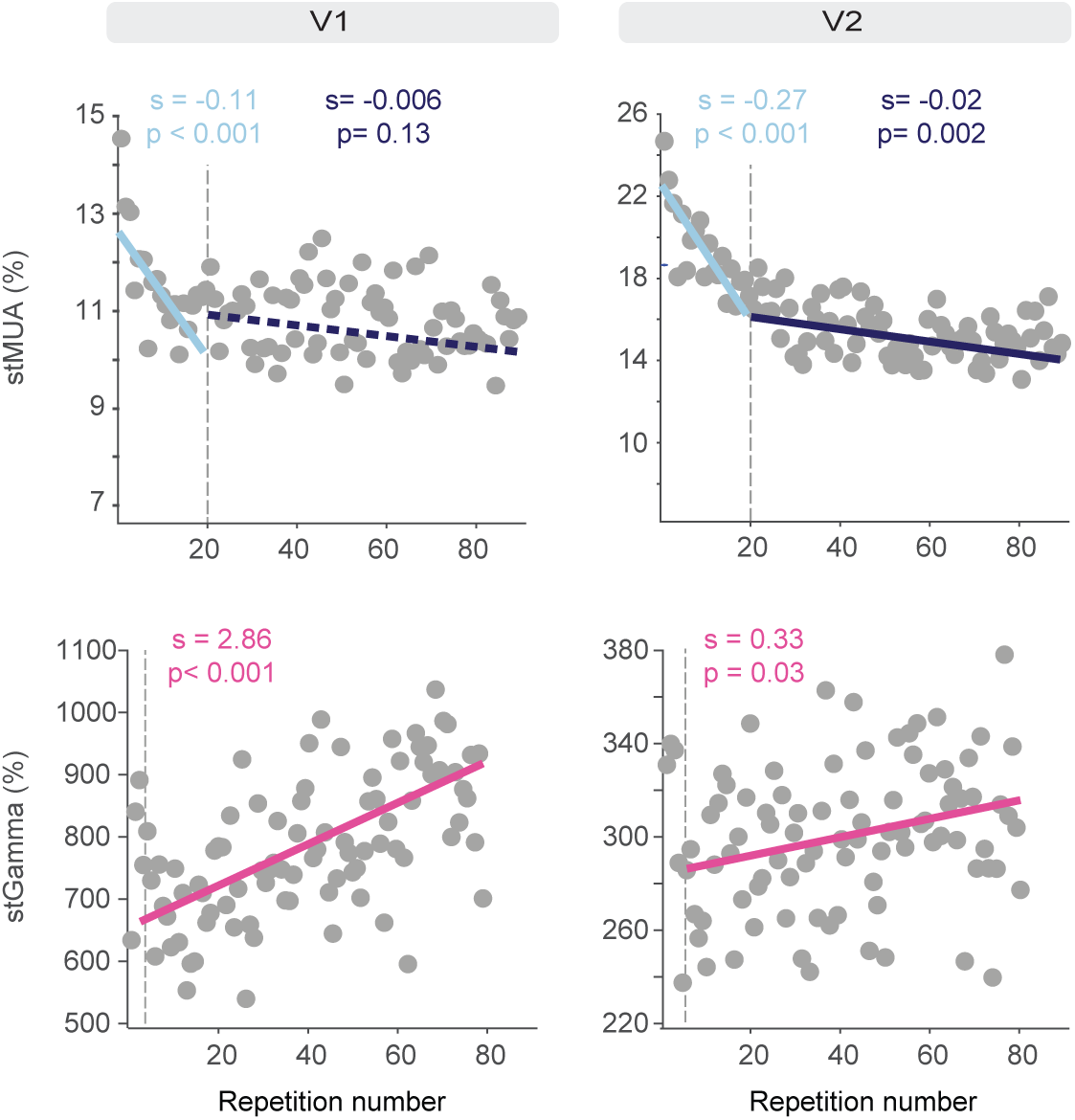
Single-trial estimates averaged over laminar compartments during FIX. stMUA (top) and stGamma (bottom) were averaged over laminar compartments of V1 (left) and V2 (right). For stMUA, the dashed vertical line indicates trial 20, and two separate regressions were performed for the Early-20 repetition group and the Remaining-70 repetition group (see Fig. 2A). For stGamma, the dashed vertical line indicates trial 7, and one regression was performed for repetitions 7-90. The slopes (s) and p-values (p) for each fit are color-coded and reported on top of each panel.

**Figure S4.**
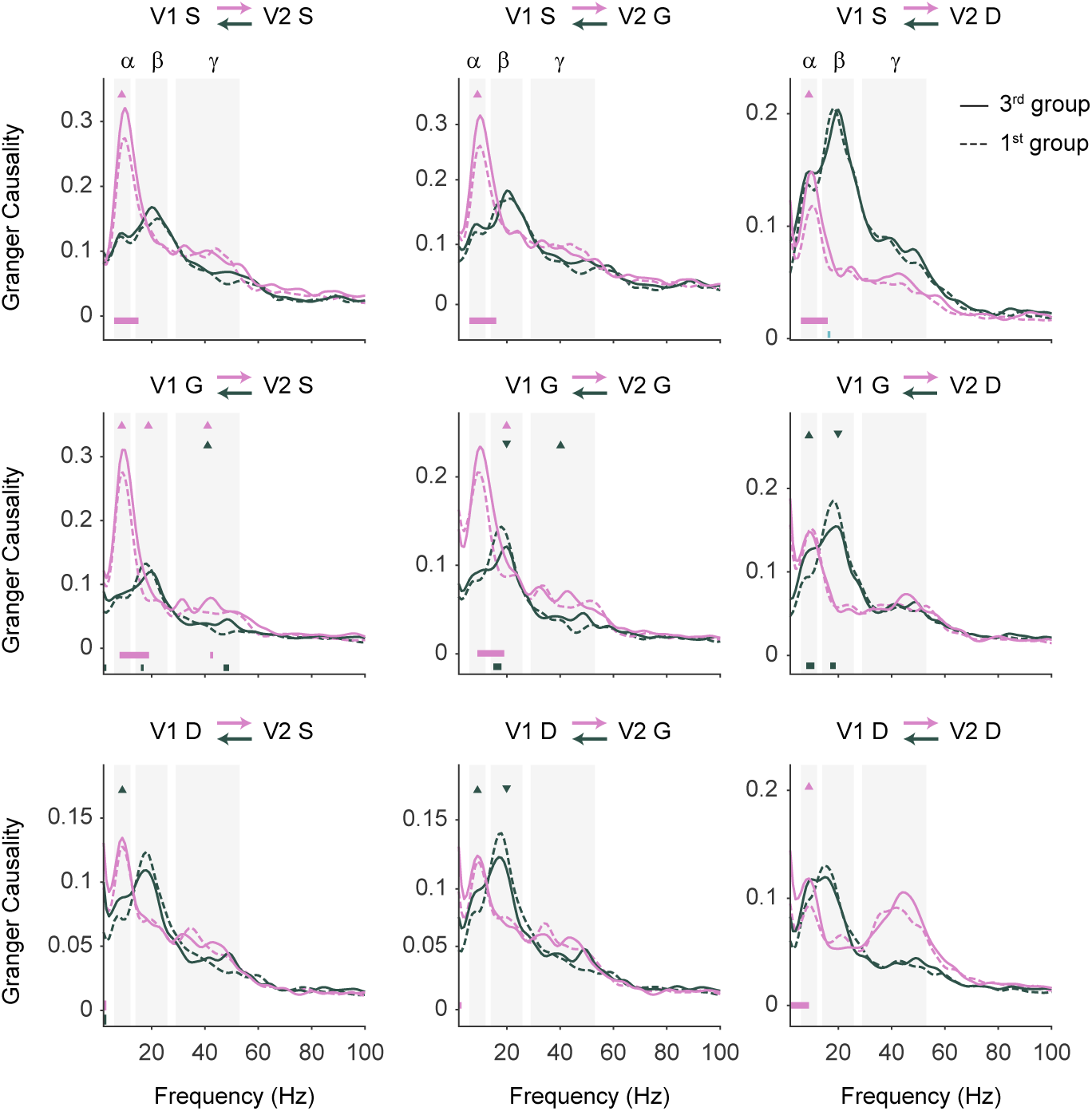
Granger Causality between laminar compartments of areas V1 and V2 during the 1^st^ repetition group (dashed line) and 3^rd^ repetition group (solid line) of FIX. Note that the RF centers of the simultaneously recorded sites in V1 and V2 (N = 9 sessions) were largely non-overlapping (see Fig. S1C-D). Different colors represent the GC direction as indicated on top of each panel. Grey rectangles show the three frequency bands that were used for statistical comparison: alpha (α), beta (β), and gamma (γ). Triangles show statistical significance per frequency band, with the triangles pointing upward for increases and downward for decreases. Horizontal lines at the bottom of the plots indicate significant differences across frequencies, color-coded for feedforward and feedback directions.

**Figure S5.**
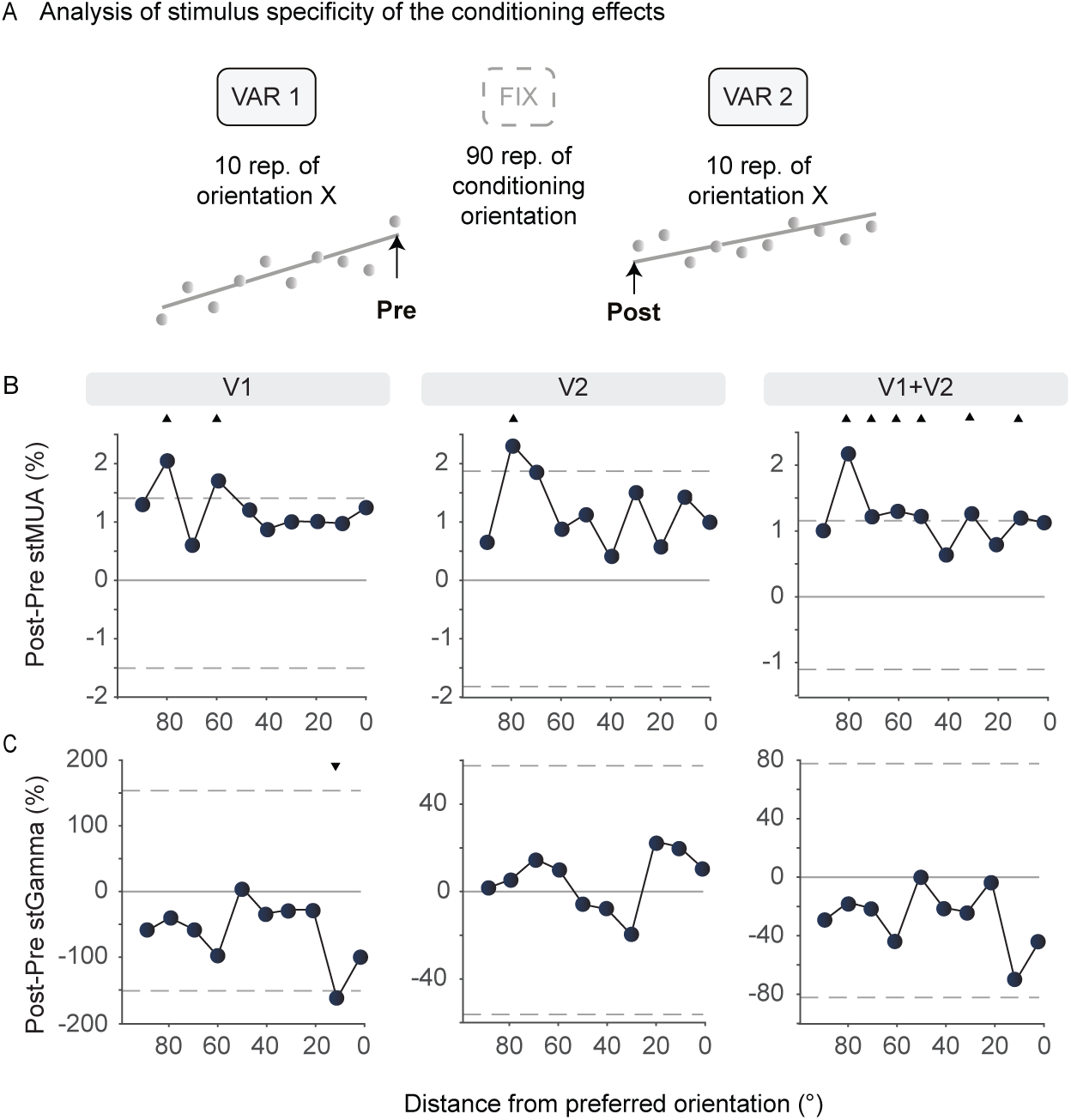
Control analysis: Repetition-related changes relative to the orientation distance from the preferred orientation. Same as Fig. 6, but as a function of the orientation distance from the preferred orientation. To ease direct comparison, the Y-axis limits are the same as in the respective panels of Fig. 6.

**Figure S6.**
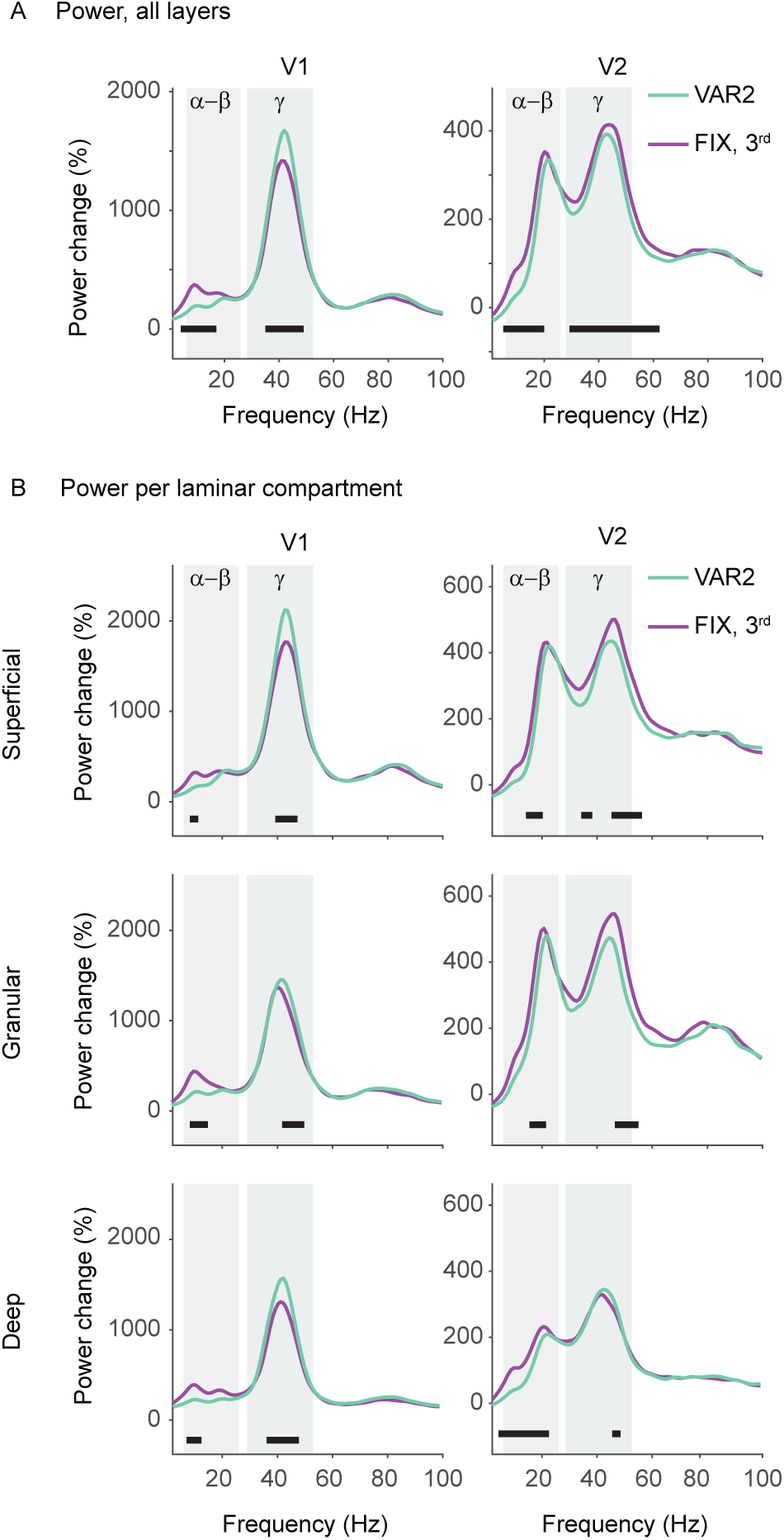
Comparison of average power between FIX and VAR2. (A) Power change relative to baseline in response to the conditioned orientation in the 3^rd^ repetition group in FIX (purple) and in VAR2 (light green), separately for V1 (left) and V2 (right). (B) Same as (A) but per laminar compartment. Black horizontal lines denote frequencies where significant differences were observed between the two conditions. Grey rectangles indicate the alpha-beta and gamma frequency bands that were used for single-trial estimates in Fig. 7. The left panel of (A) is identical to Fig. 7b and is reproduced here to facilitate the direct comparison between V1 and V2.

**Figure S7.**
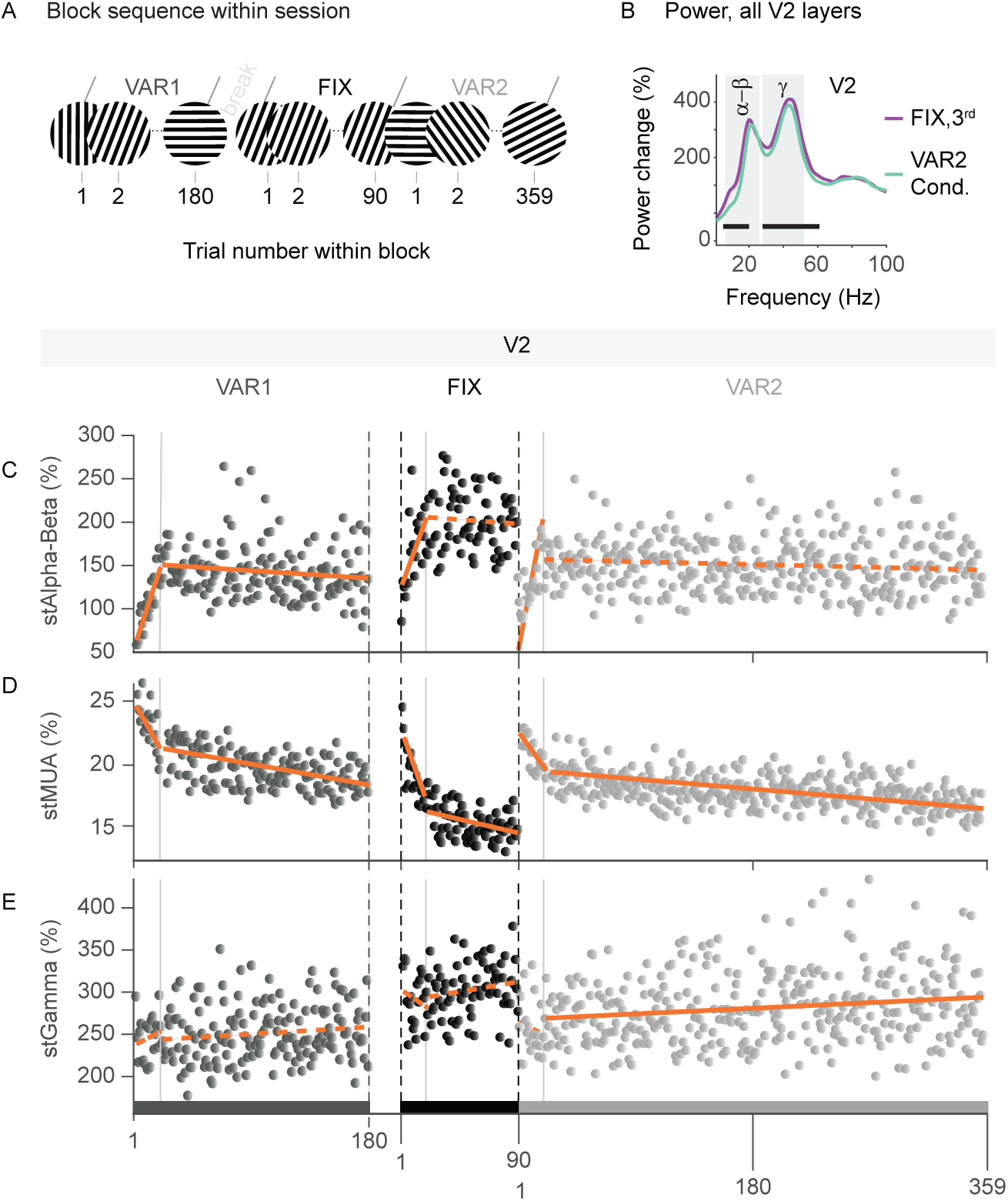
Comparison of neuronal responses in V2 across blocks. Same as Fig. 7, but for area V2. The slopes and p-values are summarized in Table S1. A statistical comparison of the regression means are presented in Table 2.

**Table S1.**
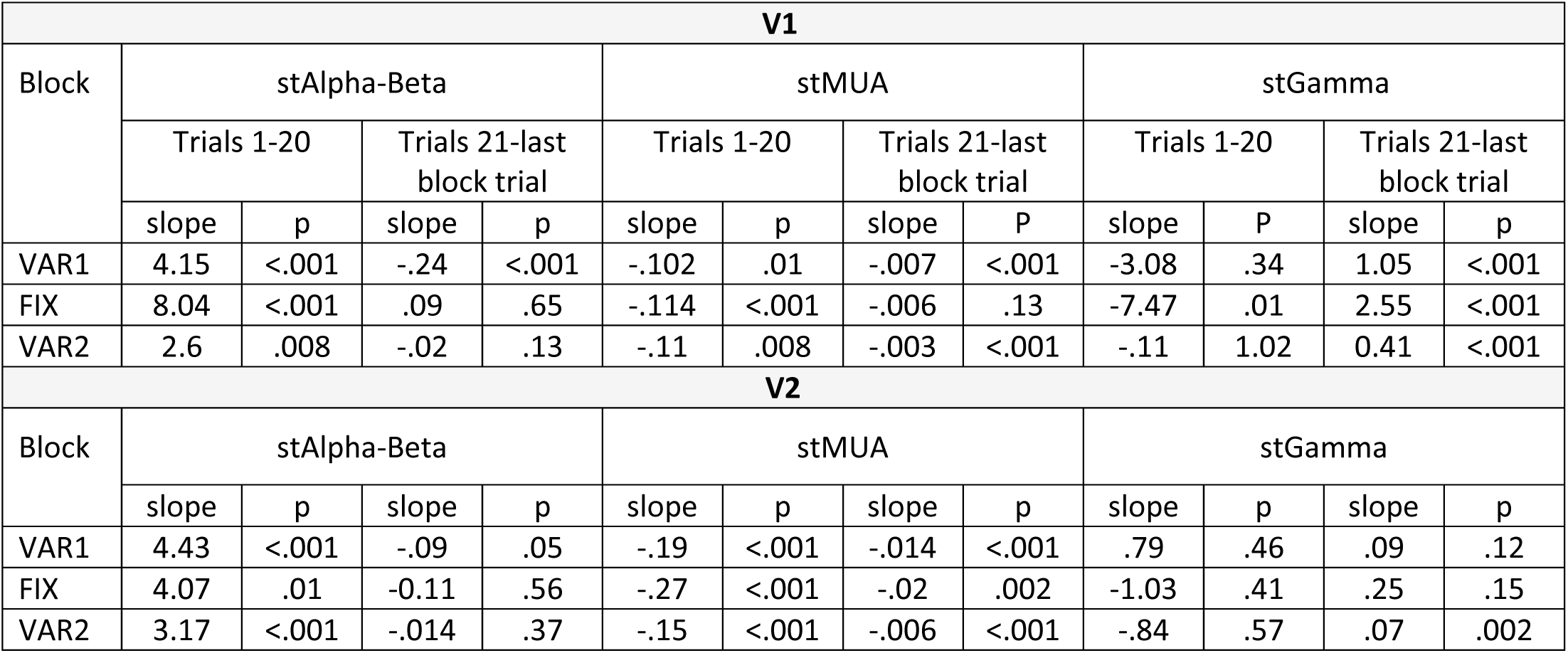
Summary of regression slopes and their p-values for Fig. 7 and Fig. S7.

| **V1** |  |  |  |  |  |  |  |  |  |  |  |  |
| --- | --- | --- | --- | --- | --- | --- | --- | --- | --- | --- | --- | --- |
| Block | stAlpha-Beta |  |  |  | stMUA |  |  |  | stGamma |  |  |  |
|  | Trials 1-20 |  | Trials 21-last block trial |  | Trials 1-20 |  | Trials 21-last block trial |  | Trials 1-20 |  | Trials 21-last block trial |  |
|  | slope | p | slope | p | slope | p | slope | P | slope | P | slope | p |
| VAR1 | 4.15 | <.001 | -.24 | <.001 | -.102 | .01 | -.007 | <.001 | -3.08 | .34 | 1.05 | <.001 |
| FIX | 8.04 | <.001 | .09 | .65 | -.114 | <.001 | -.006 | .13 | -7.47 | .01 | 2.55 | <.001 |
| VAR2 | 2.6 | .008 | -.02 | .13 | -.11 | .008 | -.003 | <.001 | -.11 | 1.02 | 0.41 | <.001 |
| **V2** |  |  |  |  |  |  |  |  |  |  |  |  |
| Block | stAlpha-Beta |  |  |  | stMUA |  |  |  | stGamma |  |  |  |
|  | slope | p | slope | p | slope | p | slope | p | slope | p | slope | p |
| VAR1 | 4.43 | <.001 | -.09 | .05 | -.19 | <.001 | -.014 | <.001 | .79 | .46 | .09 | .12 |
| FIX | 4.07 | .01 | -0.11 | .56 | -.27 | <.001 | -.02 | .002 | -1.03 | .41 | .25 | .15 |
| VAR2 | 3.17 | <.001 | -.014 | .37 | -.15 | <.001 | -.006 | <.001 | -.84 | .57 | .07 | .002 |

## CRediT author contributions

**Eleni Psarou:** Conceptualization, Methodology, Investigation, Software, Formal analysis, Visualization, Writing - Original Draft, Writing - Review & Editing. **Mohsen Parto Dezfouli:** Software, Formal analysis, Visualization, Writing - Review & Editing. **Iris Grothe:** Methodology, Investigation, Writing - Review & Editing. **Alina Peter:** Conceptualization, Writing - Review & Editing. **Rasmus Roese:** Methodology, Investigation, Writing - Review & Editing. **Pascal Fries:** Conceptualization, Methodology, Investigation, Writing - Original Draft, Writing - Review & Editing, Supervision, Project administration, Resources, Funding acquisition.

## Acknowledgements

We thank Christini Katsanevaki, Dr. Benjamin Stauch, and Dr. Yufeng Zhang for inspiring discussions and invaluable scientific input. We thank Marianne Hartmann, Julia Hoffmann, Sabrina Wallrath, and Johanna Klon-Lipok for their assistance in animal care and surgical procedures. We are grateful to Dr. Christa Tandi and Dr. Alf Theisen who provided excellent veterinary care and advice. We also thank the technical service teams of the Ernst Strüngmann Institute for their exceptional technical support. P.F. acknowledges grant support by the German Research Foundation (FR2557/1-1, FR2557/2-1, FR2557/5-1, FR2557/7-1), the European Union (HEALTH-F2-2008-200728-BrainSynch, FP7-604102-HBP) and a European Young Investigator Award.

## References

1. Hayhoe M, Ballard D. Eye movements in natural behavior. Trends Cogn Sci. 2005;9(4):188–94. Epub 2005/04/06. doi: 10.1016/j.tics.2005.02.009. PubMed PMID: 15808501.

2. Li L, Miller EK, Desimone R. The representation of stimulus familiarity in anterior inferior temporal cortex. Journal of neurophysiology. 1993;69(6):1918–29. Epub 1993/06/01. PubMed PMID: 8350131.

3. Brunet NM, Bosman CA, Vinck M, Roberts M, Oostenveld R, Desimone R, et al. Stimulus repetition modulates gamma-band synchronization in primate visual cortex. Proc Natl Acad Sci U S A. 2014;111(9):3626–31. Epub 2014/02/21. doi: 10.1073/pnas.1309714111. PubMed PMID: 24554080; PubMed Central PMCID: PMC3948273.

4. Peter A, Stauch BJ, Shapcott KA, Kouroupaki K, Schmiedt JT, Klein L, et al. Stimulus-specific plasticity of macaque V1 spike rates and gamma. Cell reports. 2021;37(10):110086. doi: 10.1016/j.celrep.2021.110086.

5. Miller EK, Li L, Desimone R. Activity of neurons in anterior inferior temporal cortex during a short-term memory task. J Neurosci. 1993;13(4):1460–78. Epub 1993/04/01. PubMed PMID: 8463829.

6. Grill-Spector K, Henson R, Martin A. Repetition and the brain: neural models of stimulus-specific effects. Trends Cogn Sci. 2006;10(1):14–23. Epub 2005/12/03. doi: 10.1016/j.tics.2005.11.006. PubMed PMID: 16321563.

7. Sawamura H, Georgieva S, Vogels R, Vanduffel W, Orban GA. Using functional magnetic resonance imaging to assess adaptation and size invariance of shape processing by humans and monkeys. J Neurosci. 2005;25(17):4294–306. Epub 2005/04/29. doi: 10.1523/jneurosci.0377-05.2005. PubMed PMID: 15858056; PubMed Central PMCID: PMCPMC6725102.

8. Stauch BJ, Peter A, Schuler H, Fries P. Stimulus-specific plasticity in human visual gamma-band activity and functional connectivity. Elife. 2021;10:e68240. Epub 2021/09/03. doi: 10.7554/eLife.68240. PubMed PMID: 34473058; PubMed Central PMCID: PMC8412931.

9. Wiggs CL, Martin A. Properties and mechanisms of perceptual priming. Current Opinion in Neurobiology. 1998;8(2):227–33. doi: 10.1016/S0959-4388(98)80144-X.

10. McMahon DB, Olson CR. Repetition suppression in monkey inferotemporal cortex: relation to behavioral priming. Journal of neurophysiology. 2007;97(5):3532–43. Epub 2007/03/09. doi: 10.1152/jn.01042.2006. PubMed PMID: 17344370.

11. Gotts SJ, Chow CC, Martin A. Repetition priming and repetition suppression: A case for enhanced efficiency through neural synchronization. Cognitive neuroscience. 2012;3(3-4):227–37. Epub 2012/11/13. doi: 10.1080/17588928.2012.670617. PubMed PMID: 23144664; PubMed Central PMCID: PMC3491809.

12. Gilbert JR, Gotts SJ, Carver FW, Martin A. Object repetition leads to local increases in the temporal coordination of neural responses. Frontiers in human neuroscience. 2010;4:30. Epub 2010/05/14. doi: 10.3389/fnhum.2010.00030. PubMed PMID: 20463867; PubMed Central PMCID: PMC2868300.

13. Rao RP, Ballard DH. Predictive coding in the visual cortex: a functional interpretation of some extra-classical receptive-field effects. Nature neuroscience. 1999;2(1):79–87. Epub 1999/04/09. doi: 10.1038/4580. PubMed PMID: 10195184.

14. Friston K. A theory of cortical responses. Philosophical transactions of the Royal Society of London Series B, Biological sciences. 2005;360(1456):815-36. Epub 2005/06/07. doi: 10.1098/rstb.2005.1622. PubMed PMID: 15937014; PubMed Central PMCID: PMCPMC1569488.

15. Mumford D. On the computational architecture of the neocortex. II. The role of cortico-cortical loops. Biological cybernetics. 1992;66(3):241–51. Epub 1992/01/01. doi: 10.1007/bf00198477. PubMed PMID: 1540675.

16. Rockland KS, Pandya DN. Laminar origins and terminations of cortical connections of the occipital lobe in the rhesus monkey. Brain research. 1979;179(1):3–20. Epub 1979/12/21. doi: 10.1016/0006-8993(79)90485-2. PubMed PMID: 116716.

17. Cragg BG. The topography of the afferent projections in the circumstriate visual cortex of the monkey studied by the Nauta method. Vision Res. 1969;9(7):733–47. Epub 1969/07/01. doi: 10.1016/0042-6989(69)90011-x. PubMed PMID: 4979024.

18. Wong-Riley M. Reciprocal connections between striate and prestriate cortex in squirrel monkey as demonstrated by combined peroxidase histochemistry and autoradiography. Brain research. 1978;147(1):159–64. doi: 10.1016/0006-8993(78)90781-3.

19. Markov NT, Vezoli J, Chameau P, Falchier A, Quilodran R, Huissoud C, et al. Anatomy of hierarchy: feedforward and feedback pathways in macaque visual cortex. The Journal of comparative neurology. 2014;522(1):225–59. Epub 2013/08/29. doi: 10.1002/cne.23458. PubMed PMID: 23983048.

20. Buffalo EA, Fries P, Landman R, Buschman TJ, Desimone R. Laminar differences in gamma and alpha coherence in the ventral stream. Proc Natl Acad Sci U S A. 2011;108(27):11262–7. Epub 2011/06/22. doi: 10.1073/pnas.1011284108. PubMed PMID: 21690410; PubMed Central PMCID: PMC3131344.

21. Bastos AM, Lundqvist M, Waite AS, Kopell N, Miller EK. Layer and rhythm specificity for predictive routing. Proc Natl Acad Sci U S A. 2020. Epub 2020/11/25. doi: 10.1073/pnas.2014868117. PubMed PMID: 33229572.

22. Xing D, Yeh CI, Burns S, Shapley RM. Laminar analysis of visually evoked activity in the primary visual cortex. Proc Natl Acad Sci U S A. 2012;109(34):13871–6. Epub 2012/08/09. doi: 10.1073/pnas.1201478109. PubMed PMID: 22872866; PubMed Central PMCID: PMC3427063.

23. Gieselmann MA, Thiele A. Stimulus dependence of directed information exchange between cortical layers in macaque V1. Elife. 2022;11. Epub 2022/03/12. doi: 10.7554/eLife.62949. PubMed PMID: 35274614; PubMed Central PMCID: PMCPMC8916775.

24. Lowet E, Roberts MJ, Peter A, Gips B, De Weerd P. A quantitative theory of gamma synchronization in macaque V1. eLife. 2017;6:e26642. doi: 10.7554/eLife.26642.

25. Drebitz E, Rausch L-P, Domingo Gil E, Kreiter AK. Three distinct gamma oscillatory networks within cortical columns in macaque monkeys’ area V1. 2024;Volume 18 - 2024. doi: 10.3389/fncir.2024.1490638.

26. Bastos AM, Vezoli J, Bosman CA, Schoffelen JM, Oostenveld R, Dowdall JR, et al. Visual areas exert feedforward and feedback influences through distinct frequency channels. Neuron. 2015;85(2):390–401. Epub 2015/01/06. doi: 10.1016/j.neuron.2014.12.018. PubMed PMID: 25556836.

27. Vezoli J, Vinck M, Bosman CA, Bastos AM, Lewis CM, Kennedy H, et al. Brain rhythms define distinct interaction networks with differential dependence on anatomy. Neuron. 2021;109(23):3862–78 e5. Epub 2021/10/22. doi: 10.1016/j.neuron.2021.09.052. PubMed PMID: 34672985; PubMed Central PMCID: PMCPMC8639786.

28. Buschman TJ, Miller EK. Top-down versus bottom-up control of attention in the prefrontal and posterior parietal cortices. Science. 2007;315(5820):1860-2. Epub 2007/03/31. doi: 10.1126/science.1138071. PubMed PMID: 17395832.

29. van Kerkoerle T, Self MW, Dagnino B, Gariel-Mathis MA, Poort J, van der Togt C, et al. Alpha and gamma oscillations characterize feedback and feedforward processing in monkey visual cortex. Proc Natl Acad Sci U S A. 2014. Epub 2014/09/11. doi: 10.1073/pnas.1402773111. PubMed PMID: 25205811.

30. Michalareas G, Vezoli J, van Pelt S, Schoffelen JM, Kennedy H, Fries P. Alpha-Beta and Gamma Rhythms Subserve Feedback and Feedforward Influences among Human Visual Cortical Areas. Neuron. 2016;89(2):384–97. Epub 2016/01/19. doi: 10.1016/j.neuron.2015.12.018. PubMed PMID: 26777277; PubMed Central PMCID: PMC4871751.

31. Psarou E, Vezoli J, Schölvinck ML, Ferracci PA, Zhang Y, Grothe I, et al. Modular, cement-free, customized headpost and connector-chamber implants for macaques. J Neurosci Methods. 2023;393:109899. Epub 2023/05/26. doi: 10.1016/j.jneumeth.2023.109899. PubMed PMID: 37230259.

32. Dowdall JR, Schmiedt JT, Stephan M, Fries P, editors. ARCADE: a modular multithreaded stimulus presentation software for the real-time control of stimuli, actions and reward during behavioral experiments. Society for Neuroscience; 2018; San Diego, CA.

33. Nandy AS, Nassi JJ, Reynolds JH. Laminar Organization of Attentional Modulation in Macaque Visual Area V4. Neuron. 2017;93(1):235–46. Epub 2016/12/19. doi: 10.1016/j.neuron.2016.11.029. PubMed PMID: 27989456; PubMed Central PMCID: PMCPMC5217483.

34. Drebitz E, Schledde B, Kreiter AK, Wegener D. Optimizing the Yield of Multi-Unit Activity by Including the Entire Spiking Activity. Frontiers in neuroscience. 2019;13:83. Epub 2019/02/28. doi: 10.3389/fnins.2019.00083. PubMed PMID: 30809117; PubMed Central PMCID: PMCPMC6379978.

35. Self MW, van Kerkoerle T, Super H, Roelfsema PR. Distinct roles of the cortical layers of area V1 in figure-ground segregation. Curr Biol. 2013;23(21):2121–9. Epub 2013/10/22. doi: 10.1016/j.cub.2013.09.013. PubMed PMID: 24139742.

36. Oostenveld R, Fries P, Maris E, Schoffelen JM. FieldTrip: Open source software for advanced analysis of MEG, EEG, and invasive electrophysiological data. Computational intelligence and neuroscience. 2011;2011:156869. Epub 2011/01/22. doi: 10.1155/2011/156869. PubMed PMID: 21253357; PubMed Central PMCID: PMC3021840.

37. Fiorani M, Azzi JC, Soares JG, Gattass R. Automatic mapping of visual cortex receptive fields: a fast and precise algorithm. J Neurosci Methods. 2014;221:112–26. Epub 2013/10/03. doi: 10.1016/j.jneumeth.2013.09.012. PubMed PMID: 24084390.

38. Schroeder CE, Mehta AD, Givre SJ. A spatiotemporal profile of visual system activation revealed by current source density analysis in the awake macaque. Cereb Cortex. 1998;8(7):575–92. Epub 1998/11/21. doi: 10.1093/cercor/8.7.575. PubMed PMID: 9823479.

39. Mitzdorf U. Properties of the evoked potential generators: current source-density analysis of visually evoked potentials in the cat cortex. The International journal of neuroscience. 1987;33(1-2):33–59. Epub 1987/03/01. doi: 10.3109/00207458708985928. PubMed PMID: 3610492.

40. Olsen T. Current source density (CSD) https://www.mathworks.com/matlabcentral/fileexchange/69399-current-source-density-csd: MATLAB Central File Exchange; 2023 [cited 2023 August 15].

41. Self MW, van Kerkoerle T, Goebel R, Roelfsema PR. Benchmarking laminar fMRI: Neuronal spiking and synaptic activity during top-down and bottom-up processing in the different layers of cortex. Neuroimage. 2019;197:806–17. Epub 2017/06/27. doi: 10.1016/j.neuroimage.2017.06.045. PubMed PMID: 28648888.

42. Rimehaug AE, Stasik AJ, Hagen E, Billeh YN, Siegle JH, Dai K, et al. Uncovering circuit mechanisms of current sinks and sources with biophysical simulations of primary visual cortex. Elife. 2023;12:e87169. Epub 2023/07/24. doi: 10.7554/eLife.87169. PubMed PMID: 37486105; PubMed Central PMCID: PMCPMC10393295.

43. Ziemba CM, Perez RK, Pai J, Kelly JG, Hallum LE, Shooner C, et al. Laminar Differences in Responses to Naturalistic Texture in Macaque V1 and V2. J Neurosci. 2019;39(49):9748–56. Epub 2019/11/02. doi: 10.1523/JNEUROSCI.1743-19.2019. PubMed PMID: 31666355; PubMed Central PMCID: PMCPMC6891061.

44. Kelly JG, Hawken MJ. Quantification of neuronal density across cortical depth using automated 3D analysis of confocal image stacks. Brain Struct Funct. 2017;222(7):3333–53. Epub 2017/03/01. doi: 10.1007/s00429-017-1382-6. PubMed PMID: 28243763; PubMed Central PMCID: PMCPMC5572544.

45. Dhamala M, Rangarajan G, Ding M. Estimating Granger causality from fourier and wavelet transforms of time series data. Physical review letters. 2008;100(1):018701. Epub 2008/02/01. PubMed PMID: 18232831.

46. Womelsdorf T, Lima B, Vinck M, Oostenveld R, Singer W, Neuenschwander S, et al. Orientation selectivity and noise correlation in awake monkey area V1 are modulated by the gamma cycle. Proc Natl Acad Sci U S A. 2012;109(11):4302–7. Epub 2012/03/01. doi: 10.1073/pnas.1114223109. PubMed PMID: 22371570; PubMed Central PMCID: PMC3306673.

47. Nichols TE, Holmes AP. Nonparametric permutation tests for functional neuroimaging: a primer with examples. Hum Brain Mapp. 2002;15(1):1–25. Epub 2001/12/18. PubMed PMID: 11747097.

48. Fries P, Maris E. What to Do If N Is Two? J Cogn Neurosci. 2022;34(7):1114–8. Epub 2022/04/26. doi: 10.1162/jocn_a_01857. PubMed PMID: 35468209.

49. Russell WMS, Burch RL. The principles of humane experimental technique: Methuen; 1959.

50. Laurens J. The statistical power of three monkeys. bioRxiv. 2022:2022.05.10.491373. doi: 10.1101/2022.05.10.491373.

51. Psarou E, Katsanevaki C, Maris E, Fries P. Would you agree if N is three? On statistical inference for small N. bioRxiv. 2024:2024.08.26.609821. doi: 10.1101/2024.08.26.609821.

52. Gilbert CD. Laminar differences in receptive field properties of cells in cat primary visual cortex. J Physiol. 1977;268(2):391–421. Epub 1977/06/01. doi: 10.1113/jphysiol.1977.sp011863. PubMed PMID: 874916; PubMed Central PMCID: PMCPMC1283670.

53. Friese U, Supp GG, Hipp JF, Engel AK, Gruber T. Oscillatory MEG gamma band activity dissociates perceptual and conceptual aspects of visual object processing: a combined repetition/conceptual priming study. Neuroimage. 2012;59(1):861–71. Epub 2011/08/13. doi: 10.1016/j.neuroimage.2011.07.073. PubMed PMID: 21835246.

54. Gruber T, Müller MM. Effects of picture repetition on induced gamma band responses, evoked potentials, and phase synchrony in the human EEG. Brain research Cognitive brain research. 2002;13(3):377–92. Epub 2002/03/29. PubMed PMID: 11919002.

55. Bastos AM, Loonis R, Kornblith S, Lundqvist M, Miller EK. Laminar recordings in frontal cortex suggest distinct layers for maintenance and control of working memory. Proc Natl Acad Sci U S A. 2018;115(5):1117–22. Epub 2018/01/18. doi: 10.1073/pnas.1710323115. PubMed PMID: 29339471; PubMed Central PMCID: PMC5798320.

56. Roberts MJ, Lowet E, Brunet NM, Ter Wal M, Tiesinga P, Fries P, et al. Robust gamma coherence between macaque V1 and V2 by dynamic frequency matching. Neuron. 2013;78(3):523–36. Epub 2013/05/15. doi: 10.1016/j.neuron.2013.03.003. PubMed PMID: 23664617.

57. Fries P, Nikolić D, Singer W. The gamma cycle. Trends in neurosciences. 2007;30(7):309-16. Epub 2007/06/09. doi: 10.1016/j.tins.2007.05.005. PubMed PMID: 17555828.

58. Vinck M, Lima B, Womelsdorf T, Oostenveld R, Singer W, Neuenschwander S, et al. Gamma-phase shifting in awake monkey visual cortex. J Neurosci. 2010;30(4):1250–7. Epub 2010/01/29. doi: 10.1523/JNEUROSCI.1623-09.2010. PubMed PMID: 20107053.

59. Atallah BV, Scanziani M. Instantaneous modulation of gamma oscillation frequency by balancing excitation with inhibition. Neuron. 2009;62(4):566–77. Epub 2009/05/30. doi: 10.1016/j.neuron.2009.04.027. PubMed PMID: 19477157; PubMed Central PMCID: PMC2702525.

60. Vinck M, Womelsdorf T, Buffalo EA, Desimone R, Fries P. Attentional modulation of cell-class-specific gamma-band synchronization in awake monkey area V4. Neuron. 2013;80(4):1077–89. Epub 2013/11/26. doi: 10.1016/j.neuron.2013.08.019. PubMed PMID: 24267656; PubMed Central PMCID: PMC3840396.

61. Hasenstaub A, Shu Y, Haider B, Kraushaar U, Duque A, McCormick DA. Inhibitory postsynaptic potentials carry synchronized frequency information in active cortical networks. Neuron. 2005;47(3):423–35. Epub 2005/08/02. doi: 10.1016/j.neuron.2005.06.016. PubMed PMID: 16055065.

62. Csicsvari J, Jamieson B, Wise KD, Buzsáki G. Mechanisms of gamma oscillations in the hippocampus of the behaving rat. Neuron. 2003;37(2):311–22. Epub 2003/01/28. PubMed PMID: 12546825.

63. Ahissar E, Vaadia E, Ahissar M, Bergman H, Arieli A, Abeles M. Dependence of cortical plasticity on correlated activity of single neurons and on behavioral context. Science (New York, NY). 1992;257(5075):1412-5. Epub 1992/09/04. doi: 10.1126/science.1529342. PubMed PMID: 1529342.

64. Caporale N, Dan Y. Spike Timing–Dependent Plasticity: A Hebbian Learning Rule. 2008;31(Volume 31, 2008):25–46. doi: 10.1146/annurev.neuro.31.060407.125639.

65. Womelsdorf T, Fries P, Mitra PP, Desimone R. Gamma-band synchronization in visual cortex predicts speed of change detection. Nature. 2006;439(7077):733-6. Epub 2005/12/24. doi: 10.1038/nature04258. PubMed PMID: 16372022.

66. Gieselmann MA, Thiele A. Comparison of spatial integration and surround suppression characteristics in spiking activity and the local field potential in macaque V1. The European journal of neuroscience. 2008;28(3):447–59. Epub 2008/08/16. doi: 10.1111/j.1460-9568.2008.06358.x. PubMed PMID: 18702717.

67. Peter A, Uran C, Klon-Lipok J, Roese R, van Stijn S, Barnes W, et al. Surface color and predictability determine contextual modulation of V1 firing and gamma oscillations. eLife. 2019;8. Epub 2019/02/05. doi: 10.7554/eLife.42101. PubMed PMID: 30714900; PubMed Central PMCID: PMC6391066.

68. Fries P, Womelsdorf T, Oostenveld R, Desimone R. The effects of visual stimulation and selective visual attention on rhythmic neuronal synchronization in macaque area V4. J Neurosci. 2008;28(18):4823–35. Epub 2008/05/02. doi: 10.1523/JNEUROSCI.4499-07.2008. PubMed PMID: 18448659.

69. Brunet N, Bosman CA, Roberts M, Oostenveld R, Womelsdorf T, De Weerd P, et al. Visual cortical gamma-band activity during free viewing of natural images. Cereb Cortex. 2015;25(4):918–26. Epub 2013/10/11. doi: 10.1093/cercor/bht280. PubMed PMID: 24108806; PubMed Central PMCID: PMC4379996.

70. Gregoriou GG, Gotts SJ, Desimone R. Cell-type-specific synchronization of neural activity in FEF with V4 during attention. Neuron. 2012;73(3):581–94. Epub 2012/02/14. doi: 10.1016/j.neuron.2011.12.019. PubMed PMID: 22325208; PubMed Central PMCID: PMC3297082.

71. Gregoriou GG, Gotts SJ, Zhou H, Desimone R. High-frequency, long-range coupling between prefrontal and visual cortex during attention. Science. 2009;324(5931):1207-10. Epub 2009/05/30. doi: 10.1126/science.1171402. PubMed PMID: 19478185; PubMed Central PMCID: PMC2849291.

72. Bosman CA, Schoffelen JM, Brunet N, Oostenveld R, Bastos AM, Womelsdorf T, et al. Attentional stimulus selection through selective synchronization between monkey visual areas. Neuron. 2012;75(5):875–88. Epub 2012/09/11. doi: 10.1016/j.neuron.2012.06.037. PubMed PMID: 22958827; PubMed Central PMCID: PMC3457649.

73. Angelucci A, Levitt JB, Walton EJ, Hupe JM, Bullier J, Lund JS. Circuits for local and global signal integration in primary visual cortex. J Neurosci. 2002;22(19):8633–46. Epub 2002/09/28. doi: 10.1523/JNEUROSCI.22-19-08633.2002. PubMed PMID: 12351737; PubMed Central PMCID: PMCPMC6757772.

74. McGuire BA, Gilbert CD, Rivlin PK, Wiesel TN. Targets of horizontal connections in macaque primary visual cortex. J Comp Neurol. 1991;305(3):370–92. Epub 1991/03/15. doi: 10.1002/cne.903050303. PubMed PMID: 1709953.

75. Stettler DD, Das A, Bennett J, Gilbert CD. Lateral connectivity and contextual interactions in macaque primary visual cortex. Neuron. 2002;36(4):739–50. Epub 2002/11/21. doi: 10.1016/s0896-6273(02)01029-2. PubMed PMID: 12441061.

76. Malach R, Amir Y, Harel M, Grinvald A. Relationship between intrinsic connections and functional architecture revealed by optical imaging and in vivo targeted biocytin injections in primate striate cortex. Proc Natl Acad Sci U S A. 1993;90(22):10469–73. Epub 1993/11/15. doi: 10.1073/pnas.90.22.10469. PubMed PMID: 8248133; PubMed Central PMCID: PMCPMC47798.

77. Das A, Gilbert CD. Long-range horizontal connections and their role in cortical reorganization revealed by optical recording of cat primary visual cortex. Nature. 1995;375(6534):780-4. Epub 1995/06/29. doi: 10.1038/375780a0. PubMed PMID: 7596409.

78. Yamahachi H, Marik SA, McManus JN, Denk W, Gilbert CD. Rapid axonal sprouting and pruning accompany functional reorganization in primary visual cortex. Neuron. 2009;64(5):719–29. Epub 2009/12/17. doi: 10.1016/j.neuron.2009.11.026. PubMed PMID: 20005827; PubMed Central PMCID: PMCPMC2818836.

79. Darian-Smith C, Gilbert CD. Axonal sprouting accompanies functional reorganization in adult cat striate cortex. Nature. 1994;368:737–40. Epub 1994/04/21. doi: 10.1038/368737a0. PubMed PMID: 8152484.

80. Gilbert CD, Li W. Adult visual cortical plasticity. Neuron. 2012;75(2):250–64. Epub 2012/07/31. doi: 10.1016/j.neuron.2012.06.030. PubMed PMID: 22841310; PubMed Central PMCID: PMCPMC3408614.

81. Vinck M, Bosman CA. More Gamma More Predictions: Gamma-Synchronization as a Key Mechanism for Efficient Integration of Classical Receptive Field Inputs with Surround Predictions. Frontiers in systems neuroscience. 2016;10:35. Epub 2016/05/21. doi: 10.3389/fnsys.2016.00035. PubMed PMID: 27199684; PubMed Central PMCID: PMC4842768.

82. Uran C, Peter A, Lazar A, Barnes W, Klon-Lipok J, Shapcott KA, et al. Predictive coding of natural images by V1 firing rates and rhythmic synchronization. Neuron. 2022;110(7):1240–57.e8. doi: 10.1016/j.neuron.2022.01.002.

83. Gray CM, König P, Engel AK, Singer W. Oscillatory responses in cat visual cortex exhibit inter-columnar synchronization which reflects global stimulus properties. Nature. 1989;338(6213):334-7. Epub 1989/03/23. doi: 10.1038/338334a0. PubMed PMID: 2922061.

84. Galuske RAW, Munk MHJ, Singer W. Relation between gamma oscillations and neuronal plasticity in the visual cortex. Proc Natl Acad Sci U S A. 2019;116(46):23317–25. Epub 2019/10/30. doi: 10.1073/pnas.1901277116. PubMed PMID: 31659040; PubMed Central PMCID: PMCPMC6859324.

85. Darian-Smith C, Gilbert CD. Topographic reorganization in the striate cortex of the adult cat and monkey is cortically mediated. J Neurosci. 1995;15(3):1631–47. Epub 1995/03/01. doi: 10.1523/jneurosci.15-03-01631.1995. PubMed PMID: 7891124; PubMed Central PMCID: PMCPMC6578152.

86. Bastos AM, Briggs F, Alitto HJ, Mangun GR, Usrey WM. Simultaneous recordings from the primary visual cortex and lateral geniculate nucleus reveal rhythmic interactions and a cortical source for gamma-band oscillations. J Neurosci. 2014;34(22):7639–44. Epub 2014/05/30. doi: 10.1523/JNEUROSCI.4216-13.2014. PubMed PMID: 24872567; PubMed Central PMCID: PMC4035524.

87. Neuenschwander S, Rosso G, Branco N, Freitag F, Tehovnik EJ, Schmidt KE, et al. On the Functional Role of Gamma Synchronization in the Retinogeniculate System of the Cat. J Neurosci. 2023;43(28):5204–20. Epub 2023/06/17. doi: 10.1523/JNEUROSCI.1550-22.2023. PubMed PMID: 37328291; PubMed Central PMCID: PMCPMC10342227.

88. Katsanevaki C, Bosman CA, Friston KJ, Fries P. Stimulus-repetition effects on macaque V1 and V4 microcircuits explain gamma-synchronization increase. 2024:2024.12.06.627165. doi: 10.1101/2024.12.06.627165 %J bioRxiv.

89. Vanni S, Hokkanen H, Werner F, Angelucci A. Anatomy and Physiology of Macaque Visual Cortical Areas V1, V2, and V5/MT: Bases for Biologically Realistic Models. Cereb Cortex. 2020;30(6):3483–517. Epub 2020/01/04. doi: 10.1093/cercor/bhz322. PubMed PMID: 31897474; PubMed Central PMCID: PMCPMC7233004.

90. Markov NT, Ercsey-Ravasz MM, Ribeiro Gomes AR, Lamy C, Magrou L, Vezoli J, et al. A weighted and directed interareal connectivity matrix for macaque cerebral cortex. Cereb Cortex. 2014;24(1):17–36. Epub 2012/09/27. doi: 10.1093/cercor/bhs270. PubMed PMID: 23010748; PubMed Central PMCID: PMC3862262.

91. Hartmann TS, Raja S, Lomber SG, Born RT. Cortico-cortical feedback from V2 exerts a powerful influence over the visually evoked local field potential and associated spike timing in V1. bioRxiv. 2019:792010. doi: 10.1101/792010 %J bioRxiv.

92. Nurminen L, Merlin S, Bijanzadeh M, Federer F, Angelucci A. Top-down feedback controls spatial summation and response amplitude in primate visual cortex. Nature communications. 2018;9(1):2281. Epub 2018/06/13. doi: 10.1038/s41467-018-04500-5. PubMed PMID: 29892057; PubMed Central PMCID: PMCPMC5995810.

93. Friston K. Predictive coding, precision and synchrony. Cognitive neuroscience. 2012;3(3-4):238-9. Epub 2012/09/01. doi: 10.1080/17588928.2012.691277. PubMed PMID: 24171746.

94. Fries P, Reynolds JH, Rorie AE, Desimone R. Modulation of Oscillatory Neuronal Synchronization by Selective Visual Attention. 2001;291(5508):1560-3. doi: doi:10.1126/science.1055465.

95. Grothe I, Neitzel SD, Mandon S, Kreiter AK. Switching Neuronal Inputs by Differential Modulations of Gamma-Band Phase-Coherence. 2012;32(46):16172–80. doi: 10.1523/JNEUROSCI.0890-12.2012 %J The Journal of Neuroscience.

96. Mayer A, Schwiedrzik CM, Wibral M, Singer W, Melloni L. Expecting to See a Letter: Alpha Oscillations as Carriers of Top-Down Sensory Predictions. Cereb Cortex. 2016;26(7):3146–60. Epub 2015/07/05. doi: 10.1093/cercor/bhv146. PubMed PMID: 26142463.

97. Chao ZC, Takaura K, Wang L, Fujii N, Dehaene S. Large-Scale Cortical Networks for Hierarchical Prediction and Prediction Error in the Primate Brain. Neuron. 2018;100(5):1252–66 e3. Epub 2018/11/30. doi: 10.1016/j.neuron.2018.10.004. PubMed PMID: 30482692.

98. Feldman H, Friston KJ. Attention, uncertainty, and free-energy. Frontiers in human neuroscience. 2010;4:1–23. Epub 2010/12/17. doi: 10.3389/fnhum.2010.00215. PubMed PMID: 21160551; PubMed Central PMCID: PMCPMC3001758.

99. Brown HR, Friston KJ. Dynamic causal modelling of precision and synaptic gain in visual perception - an EEG study. Neuroimage. 2012;63(1):223–31. Epub 2012/07/04. doi: 10.1016/j.neuroimage.2012.06.044. PubMed PMID: 22750569; PubMed Central PMCID: PMCPMC3438451.

100. Fries P. Rhythms for Cognition: Communication through Coherence. Neuron. 2015;88(1):220–35. Epub 2015/10/09. doi: 10.1016/j.neuron.2015.09.034. PubMed PMID: 26447583; PubMed Central PMCID: PMC4605134.

